# Visual LLM-guided consensus spatial domain detection with L-STAR

**DOI:** 10.64898/2026.08.25.747158

**Authors:** Changyue Zhao, Zhicheng Ji

## Abstract

Spatial domain detection is a central task in spatial transcriptomics, yet existing methods exhibit highly variable performance across datasets. We introduce L-STAR, a visual LLM-guided, consensus-based framework that leverages the visual reasoning capacity of large language models to adaptively rank and integrate spatial domain detection methods. L-STAR achieves robust and consistently improved performance, outperforming single spatial domain detection methods across diverse datasets.

## Introduction

Spatial transcriptomics (ST) is a rapidly evolving class of technologies that enables gene expression profiling while preserving the spatial context of cells^1,2^. By measuring transcripts alongside their physical coordinates, ST makes it possible to study how cellular states, developmental programs, and pathological processes are organized in space, revealing biological patterns that are invisible to dissociated single-cell sequencing^3,4^. A central computational task in ST analysis is spatial domain detection, which aims to identify regions of tissue that share coherent molecular and functional characteristics. Existing spatial domain detection methods leverage a variety of modeling strategies, such as Bayesian statistical methods in BayesSpace^5^, graph-based methods in GraphST^6^, and deep learning frameworks in SpaGCN^7^ and SpaCell^8^, to integrate gene expression with spatial proximity. These approaches have been widely used to delineate anatomical layers, tissue niches, and tumor microenvironmental structures, providing a foundation for understanding spatial organization across diverse biological systems.

A recent benchmarking study spanning multiple ST platforms, tissue types, and evaluation criteria demonstrated that the performance of existing spatial domain detection methods is highly inconsistent across datasets and experimental settings, with no single method consistently outperforming others across all scenarios^9^. Method performance was shown to be highly sensitive to factors such as tissue complexity, spatial resolution, noise level, and the underlying biological structure, with different methods excelling in different settings and failing in others. As a result, achieving satisfactory performance in practice often requires careful, case-by-case selection of methods. While an ideal strategy would be to choose the optimal method for each specific dataset and biological question, doing so typically demands substantial time and human expertise, posing a major barrier to scalable and routine analysis of ST data.

Large language models (LLMs) have been shown to be broadly applicable in biomedical research^10–12^. We previously demonstrated that GPT-4 can perform cell type annotation in single-cell studies^13^, that large multimodal models achieve strong performance in classifying diverse types of biomedical images^14^, and that LLMs can optimize data preprocessing in single-cell omics^15^. Together, these findings indicate that LLMs possess substantial domain knowledge in biomedical research, exhibit visual reasoning capabilities for image analysis, and demonstrate advanced reasoning abilities for handling complex analytical tasks. Based on these observations, we hypothesize that LLMs may be capable of automatically identifying the most suitable spatial domain detection method for a given dataset, by selecting the spatial domain visualization that best aligns with the underlying tissue histology from a set of candidate results. If successful, this approach could reduce the time and expertise required for method comparison while improving the robustness and reproducibility of spatial transcriptomics analysis.

## Results

To test this hypothesis, we conducted an experiment in which GPT-5, the latest LLM developed by OpenAI, was provided with a set of three images representing a brain tissue histology image and the spatial domains identified by two different methods overlaid on the same histology image (Figure 1). These images and the corresponding performance metrics of the spatial domain detection methods were obtained from the benchmarking study^9^. In this example, the first spatial domain detection method achieved superior performance, yielding a higher adjusted Rand index (ARI) when compared with expert-annotated ground truth spatial regions. When asked to determine which method performed better, without access to the ARI values or ground truth annotations, GPT-5 correctly selected the first method. This result demonstrates the feasibility of using GPT-5 to automatically compare spatial domain detection methods and to identify an optimal method in a dataset-specific manner.

**Figure 1.**
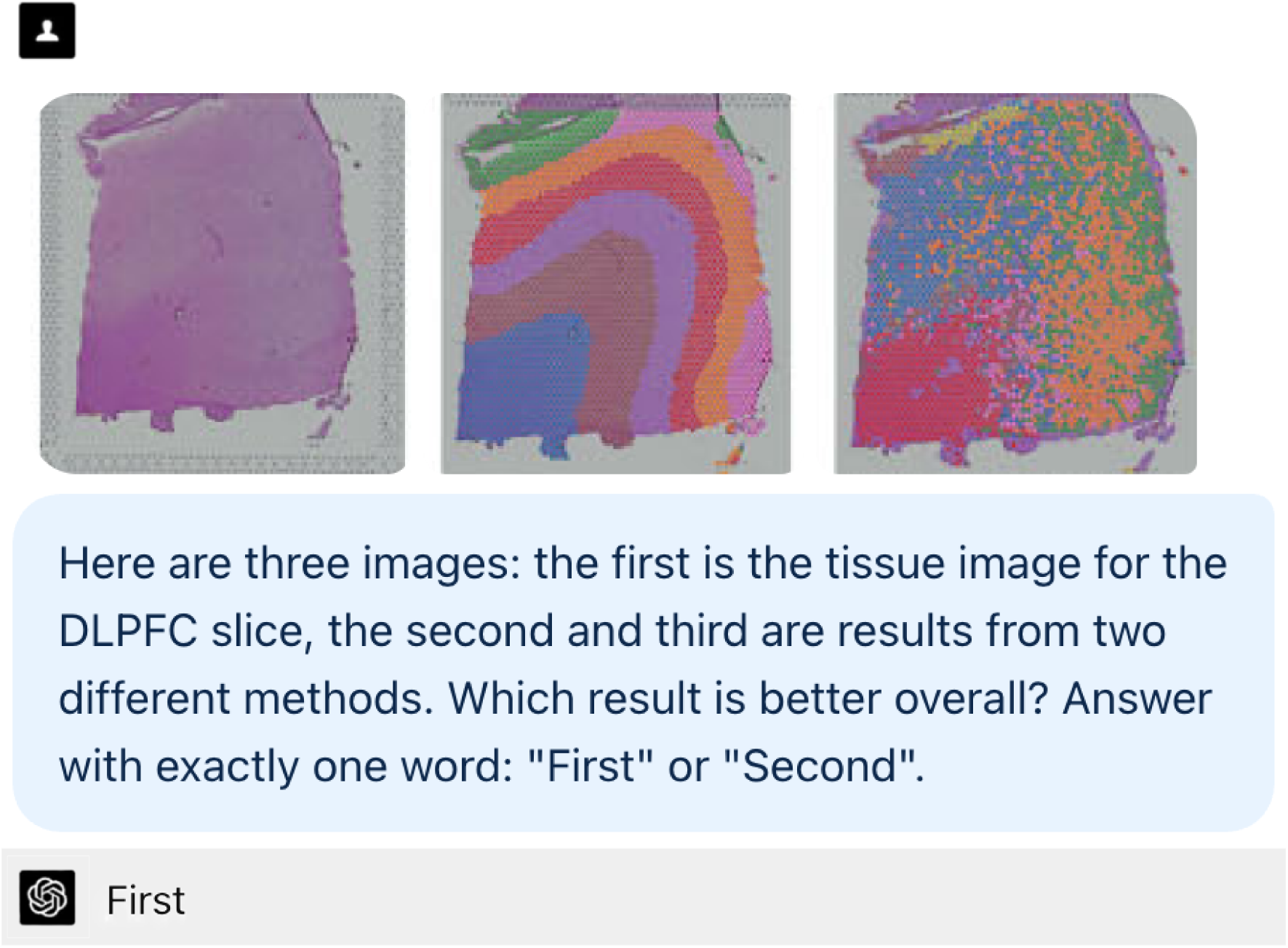
An example illustrating how an LLM visually compares spatial-domain detection results from two methods and selects the method with superior performance.

Based on this finding, we developed L-STAR, a visual **L**LM**-**guided consensus **s**pa**t**ial dom**a**in detection framework for spatial t**r**anscriptomics (Figure 2), leveraging the visual reasoning ability of contemporary LLMs. L-STAR consists of three main steps. The input to L-STAR consists of spatial domains inferred by multiple single spatial domain detection methods, such as GraphST and SpaGCN, together with the spatial coordinates of cells or spatial spots. In the first step, L-STAR generates visualizations of spatial domains based on these inputs, and GPT-5 is used to perform pairwise comparisons across all possible pairs of spatial domain detection methods. In the second step, methods are ranked according to the aggregated pairwise comparison results, and the top five methods are selected. In the third step, a consensus spatial domain is constructed by aggregating the spatial domains identified by these top-performing methods. Consensus aggregation restricted to top-performing models has previously been applied in LLM-based single-cell analysis^14^. In additional analyses, we found that L-STAR achieves its best performance when using GPT-5 as the underlying LLM, compared with other models such as Gemini and Claude (Supplementary Figure S1), and when aggregating the results from the top five methods rather than using a different number of methods (Supplementary Figure S2).

**Figure 2.**
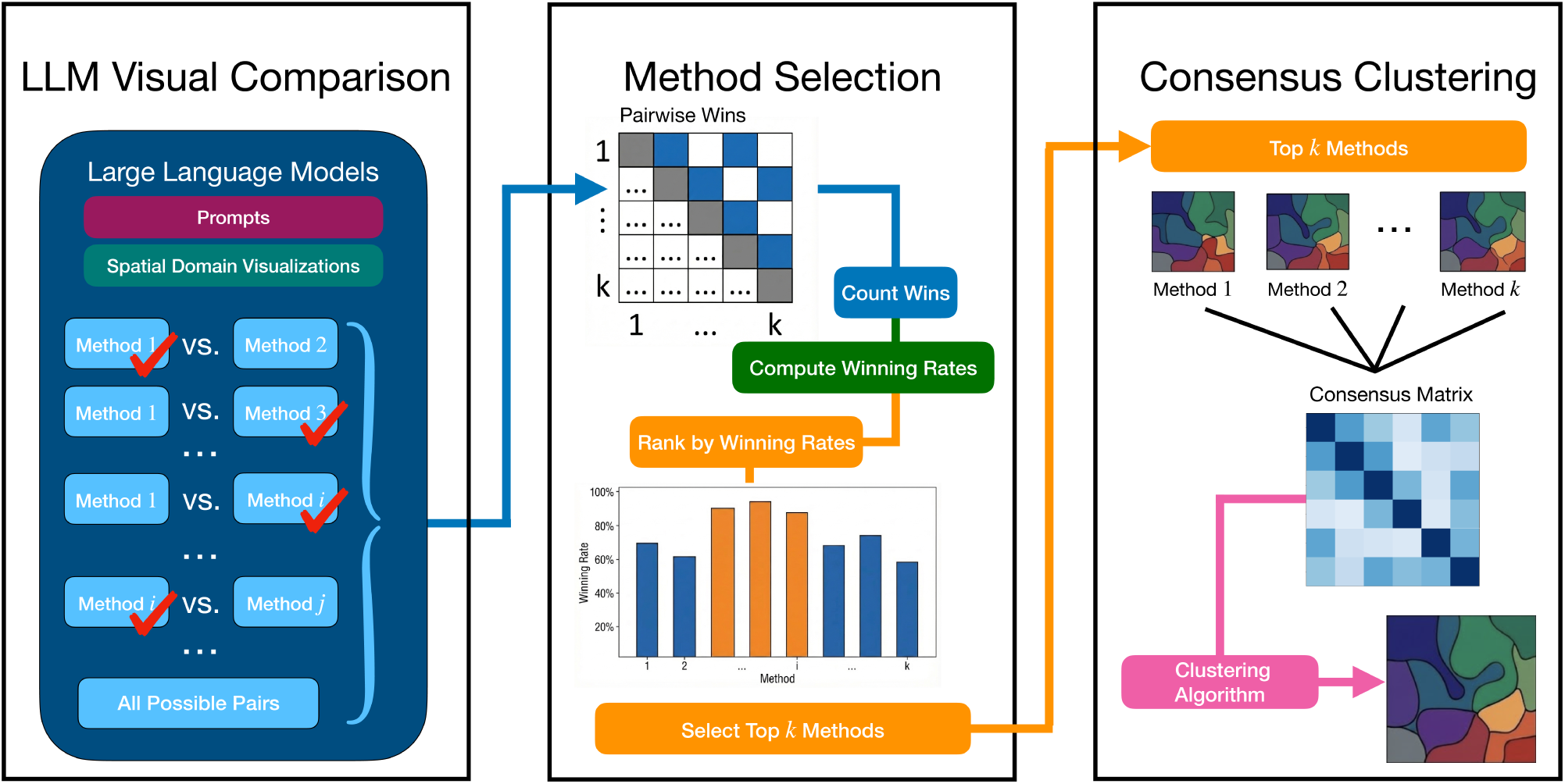
A schematic of the L-STAR workflow, comprising LLM-based pairwise comparisons of single methods, selection of top-performing methods based on winning rates, and consensus clustering to generate the final spatial domain assignments.

We first evaluated GPT-5’s ability to prioritize high-performing spatial domain detection methods using six ST datasets from the benchmark study^9^. Note that the benchmark study was published after the knowledge cutoff of GPT-5 used in this study, making data leakage unlikely. Each method’s performance was quantified by the adjusted Rand index (ARI) based on agreement between computationally derived spatial domains and expert-annotated regions. GPT-5 consistently placed the highest-ARI method within the top five across all datasets, performing substantially better than random ranking (Figure 3a). The same trend was observed using Adjusted Mutual Information (AMI) and Normalized Mutual Information (NMI) (Supplementary Figure S3a,b). These results indicate that GPT-5 can reliably identify the best method as one of the top candidates, although it may not always select it as the single top choice. This motivates the consensus strategy in L-STAR, which aggregates results from the top five methods to achieve stable performance. While LLM outputs are inherently non-deterministic, the rankings produced by GPT-5 are highly consistent across two independent runs (Supplementary Figure S4), demonstrating the robustness and reproducibility of the LLM-based ranking procedure.

**Figure 3.**
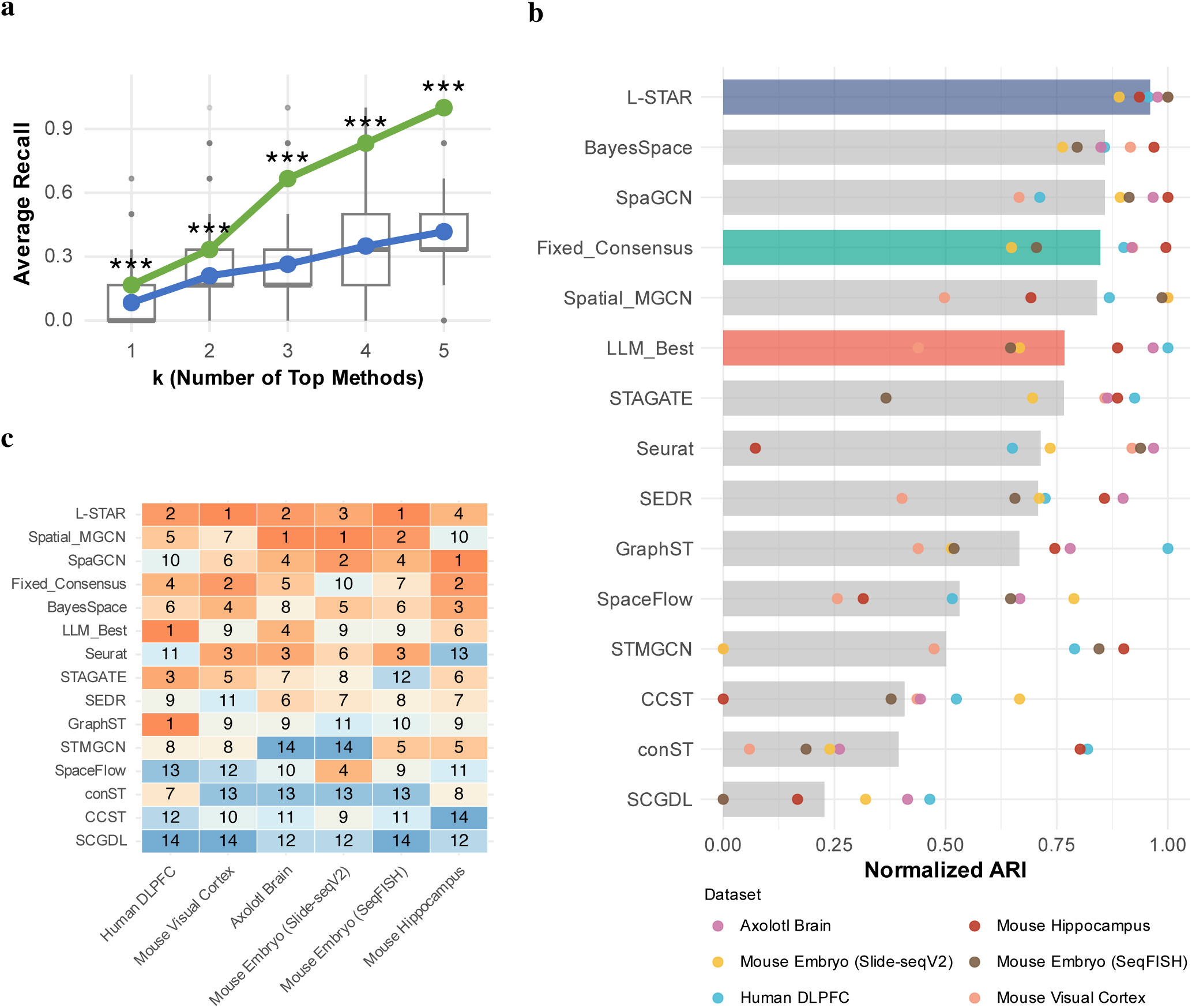
Performance of L-STAR based on the adjusted Rand index (ARI). **a**, Average recall (y-axis) as a function of the number of top-ranked methods (x-axis). Average recall is defined as the proportion of datasets for which the method achieving the highest ARI is included among the top-*k* selected methods. The green line represents the average recall when methods are ranked by GPT-5. Grey boxplots show the distribution of average recall obtained after randomly shuffling the method rankings for 5000 iterations. The blue line represents the mean recall across the 5,000 random permutations. For each fixed value of *k*, a one-sample *t*-test was used to compare the average recall from GPT-5 with that from random shuffling. *** denotes *p <* 0.001. **b**, Normalized ARI of each method. Bars represent the mean normalized ARI across datasets, while individual points indicate the normalized ARI for each dataset. “Fixed_Consensus” represents consensus clustering applied to a fixed set of methods without LLM-based selection, whereas “LLM_Best” denotes the single best method selected by GPT-5. Methods are ordered in decreasing order of their averaged normalized ARI across datasets. **c**, Ranking of each method within each dataset based on normalized ARI. Numbers and colors indicate the ranking values. Methods are ordered in increasing order of their averaged ranking across datasets.

We next compared L-STAR with several competing approaches, including single spatial domain detection methods, the highest ranked method selected by GPT-5, and consensus clustering using a fixed set of methods recommended by the benchmark study^9^. L-STAR achieved the highest average normalized ARI of 0.96 and substantially outperformed the second-best method, BayesSpace, which reached 0.86 (Figure 3b). L-STAR also ranked first under AMI and NMI (Supplementary Figure S3c,d). It also showed strong robustness, ranking within the top three in most scenarios (Figure 3c; Supplementary Figure S3e,f). These results indicate that combining GPT-5’s adaptive, dataset-specific method selection with consensus clustering offers a clear advantage, enabling L-STAR to generalize reliably across diverse tissue types. In comparison, single-method strategies, LLM-guided adaptive selection without consensus clustering, and consensus clustering without LLM-guided adaptive selection each fail in at least one setting.

Figures 4 and 5 further illustrate L-STAR’s ability to adapt across tissue types compared with single spatial domain detection methods using two representative real datasets. In the human DLPFC dataset (Figure 4), L-STAR achieves performance comparable to GraphST, the best-performing single method, and accurately recovers spatial domains consistent with the annotated cortical layers. In contrast, Spatial_MGCN exhibits clear deviations from the ground truth, failing to identify layer 2 in the first zoomed-in region and breaking cell-type homogeneity within the white matter in the second zoomed-in region. In the mouse embryo (SeqFISH) dataset, the relative performance pattern shifts (Figure 5). L-STAR again attains performance comparable to the best single method, Spatial_MGCN, whereas GraphST shows systematic discrepancies from the ground truth across both zoomed-in regions. In the first region, GraphST is dominated by a single Forebrain/Midbrain/Hindbrain domain and largely misses smaller yet biologically meaningful structures, including cranial mesoderm and definitive endoderm, resulting in an overly homogeneous assignment. In the second region, GraphST further collapses its output into a single dominant neural crest domain, thereby failing to recover the heterogeneous mixture of tissue components present in the ground truth, including endothelium, haematoendothelial progenitors, and surface ectoderm.

**Figure 4.**
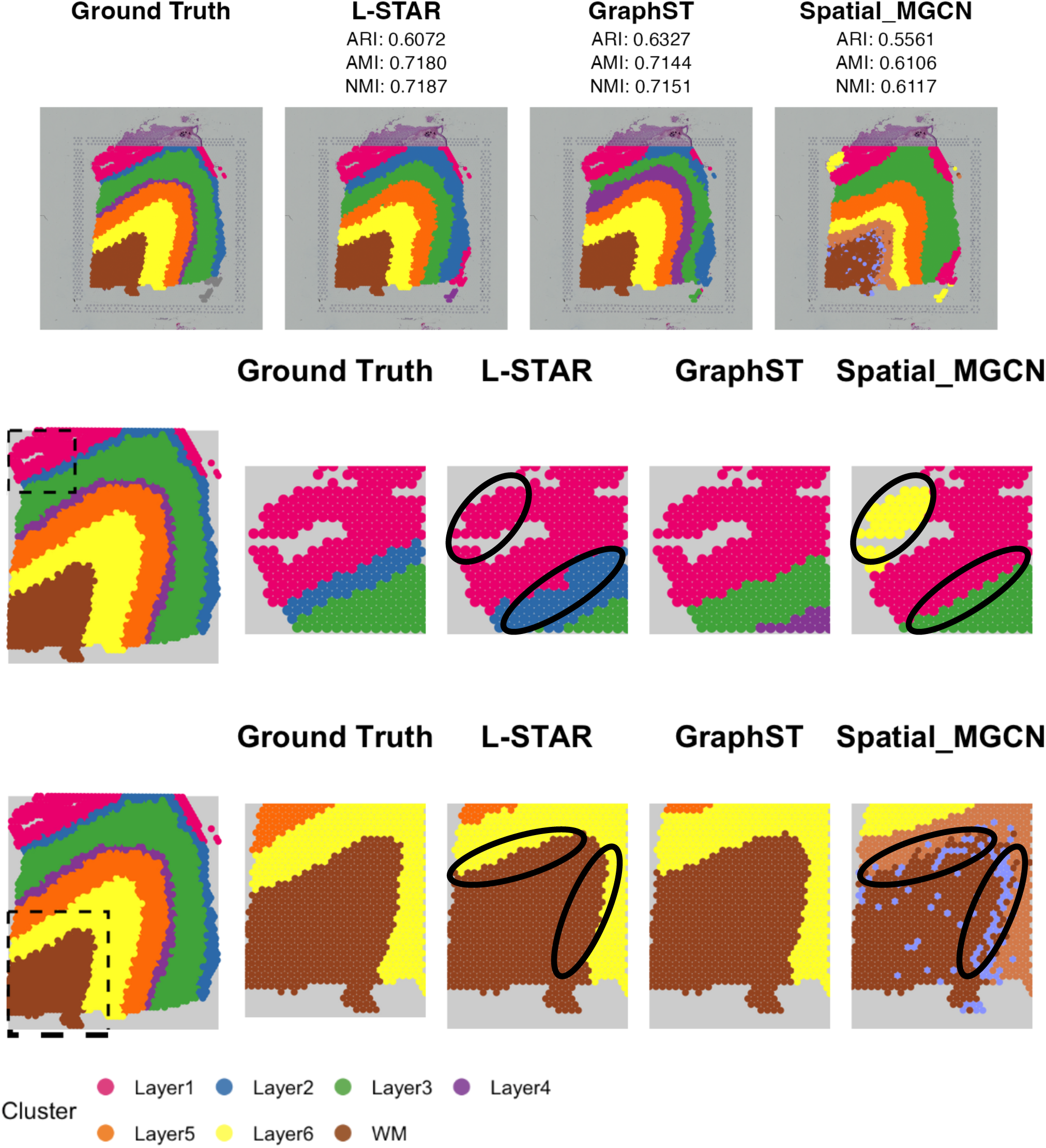
Spatial domain visualizations of the human DLPFC dataset, showing the whole-tissue view with original ARI, AMI, and NMI values (top row) and zoomed-in views (bottom two rows). Regions of interest are circled in black in the zoomed-in views.

**Figure 5.**
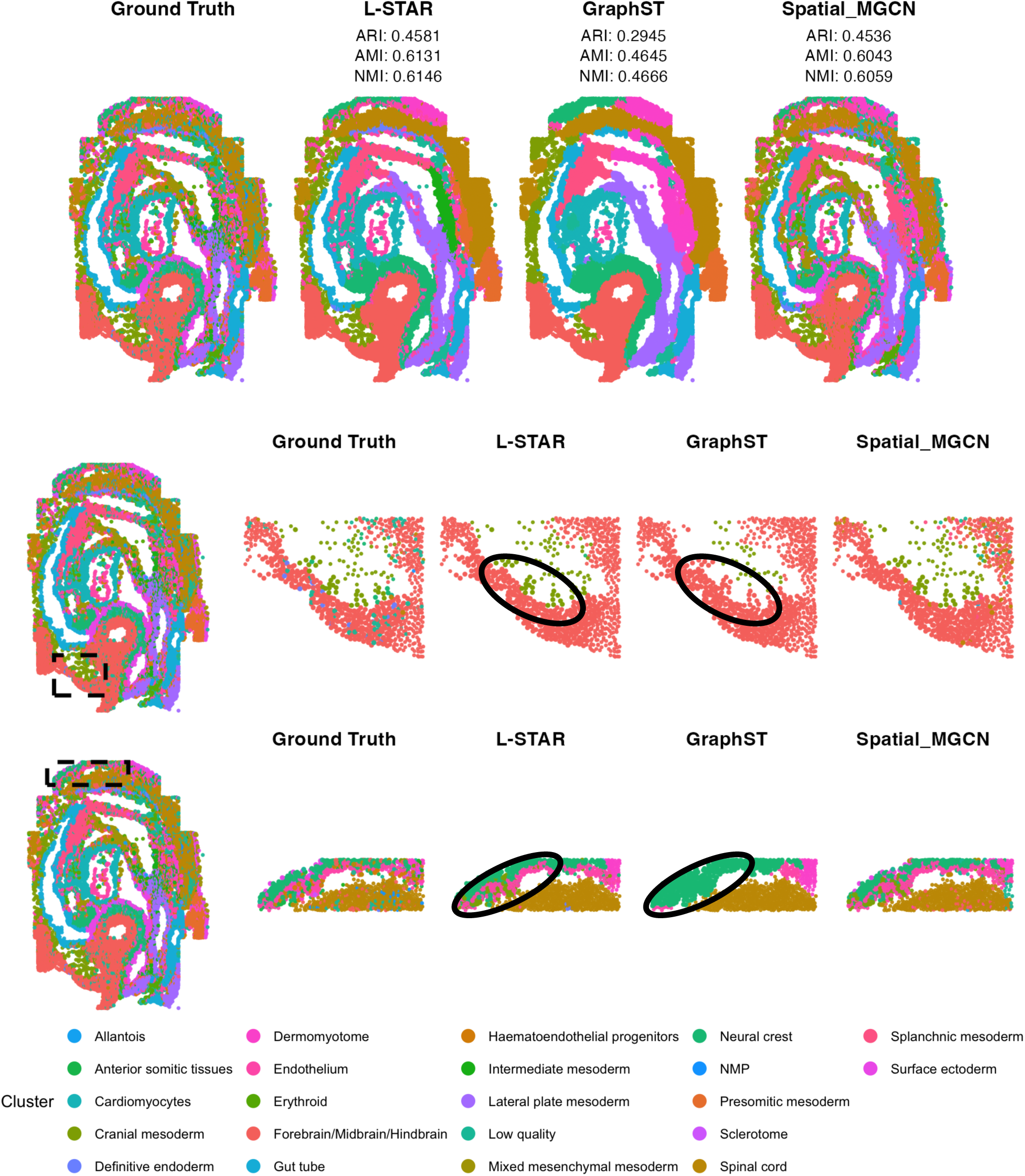
Spatial domain visualizations of the mouse embryo (SeqFISH) dataset, showing the whole-tissue view with original ARI, AMI, and NMI values (top row) and zoomed-in regions (bottom two rows). Regions of interest are circled in black in the zoomed-in views.

Beyond accuracy, the use of LLMs endows L-STAR with an interpretable decision-making process that allows users to understand the rationale underlying spatial domain detection, a capability not provided by existing methods. Figure 6a and Supplementary Figure S5 summarize the keywords extracted from the LLM-generated outputs during the pairwise comparison stage of L-STAR. These keywords capture meaningful, dataset-specific biological features. For example, in the human DLPFC dataset, terms such as “white matter” are strongly enriched relative to other slices. In contrast, in the mouse hippocampus dataset, the term “CA” (cornu ammonis) has a higher proportion of occurrence, while it is absent from other datasets and slices, reflecting hippocampus-specific anatomical structure. To provide a more direct domain-level interpretation, we developed an LLM-based post-hoc annotation module that assigns a biological name to each spatial domain identified by L-STAR. To quantitatively evaluate these annotations, we compared each LLM-generated domain name with the corresponding manual annotation curated from the original studies using the cosine similarity of their vector embeddings, and compared the observed similarities with a null distribution generated by permuting the manual annotations within each dataset. Across many spatial domains, the L-STAR annotations showed high semantic similarity to the manual annotations, and the overall similarity distribution was significantly higher than the null distribution (Figure 6b). These results indicate that the L-STAR annotations are biologically meaningful and can capture informative characteristics of the identified spatial domains. Figure 6c further illustrates an example from the DLPFC dataset, where the manual and L-STAR annotations show substantial agreement. A complete list of the manual and L-STAR annotations across the evaluated datasets is provided in Supplementary Table S1.

**Figure 6.**
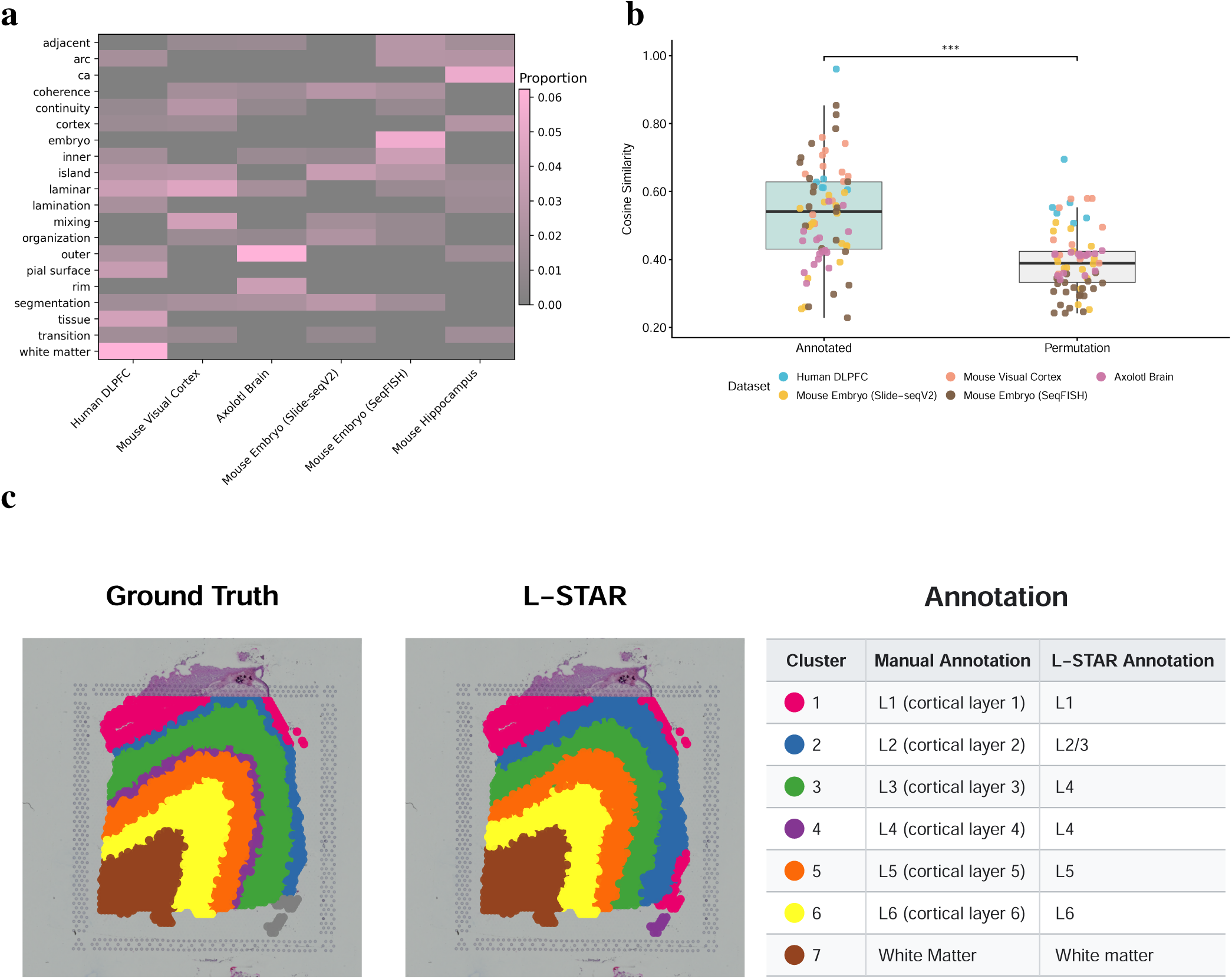
Interpretation and semantic evaluation of L-STAR outputs. **a**, Heatmap of occurrence proportions for the 20 keywords (y-axis) with the highest overall occurrence proportions across datasets (x-axis). **b**, Domain-level semantic concordance between LLM-generated and manual annotations. “Annotated” represents the observed cosine similarity, whereas “Permutation” represents each domain’s mean similarity across 10,000 within-dataset permutations of the manual annotations. Points represent individual domains and are colored by dataset. Boxes indicate the median and interquartile range, with whiskers extending to 1.5 times the interquartile range. Statistical significance was assessed using a one-sided empirical permutation test. *** denotes *p <* 0.001. **c**, Spatial domain visualizations and corresponding manual and L-STAR annotations for the human DLPFC dataset. For visualization, each manually annotated domain was matched to the L-STAR domain with the largest spot overlap, with ties resolved by the smaller L-STAR domain identifier. The two maps were colored consistently according to these overlap-based matches.

We finally quantified the computational cost and runtime of L-STAR. In our evaluation using the GPT-5 API on the six datasets, a single pairwise comparison cost an average of 0.0112 USD and required 23.15 seconds. At the dataset level, the default L-STAR pipeline cost an average of 3.70 USD and required 2.12 hours of sequential runtime per dataset. To improve scalability, we additionally implemented a cost-effective mode that compares all candidate methods simultaneously, reducing the average cost and runtime to 0.29 USD and 0.10 hours per dataset, respectively (Supplementary Figure S6). The results remained competitive with individual spatial-domain detection methods and the fixed-consensus strategy, although the default pairwise mode provided stronger overall performance in our evaluation (Supplementary Figure S7).

## Discussion

In summary, we developed L-STAR, a visual LLM-guided, consensus-based framework for spatial domain detection. L-STAR consistently achieves improved performance and adaptability across diverse datasets compared with single spatial domain detection methods, demonstrating the unique advantage of integrating the adaptive reasoning of LLMs with established computational approaches.

Because L-STAR evaluates spatial domains through their visual representations, we examined whether its performance is sensitive to the color palette used for domain visualization. Across five global hue transformations, LLM-based assessments and downstream consensus performance were generally consistent with those obtained using the default palette in most cases, indicating that L-STAR is relatively robust to reasonable changes in color palette (Supplementary Figures S8 and S9). Nevertheless, palette choice was not completely neutral, as the exact method rankings and selections could vary, and the default palette achieved the best overall ARI, AMI, and NMI across the six datasets. Removing chromatic information through grayscale also reduced the correspondence between LLM-based rankings and ground-truth performance in most cases, suggesting that color information contributes to effective visual assessment. To reduce ambiguity caused by visually similar colors assigned to adjacent domains, the default L-STAR visualization pipeline incorporates Palo^15^, a spatially aware palette optimization method that preferentially assigns distinct colors to neighboring domains. Together, these results indicate that L-STAR is generally robust to color-palette variation while supporting the use of the current standardized, spatially optimized color representation.

The default L-STAR pipeline uses exhaustive pairwise comparisons, and its number of LLM inferences therefore increases quadratically with the number of candidate methods, potentially limiting scalability for large candidate pools or cohorts. To address this limitation, L-STAR provides an optional cost-effective mode that ranks all candidate methods simultaneously, reducing the number of LLM calls from 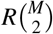 to *R*. Across the evaluated datasets, this strategy reduced both API cost and sequential runtime by more than an order of magnitude while retaining competitive downstream consensus performance (Supplementary Figures S6 and S7). These results illustrate a performance-cost tradeoff. The default pairwise mode is preferable when performance is prioritized, whereas the all-wise mode provides a scalable alternative for large candidate pools or exploratory analyses with limited computational budgets.

Recent studies have shown that LLM-based comparative judgments can exhibit position and ordering biases^16,17^, making presentation order a potential source of variability in L-STAR. Although method identities were not provided to the LLM, the default pipeline used a fixed presentation orientation for each method pair. In a sensitivity analysis using 30 independently randomized presentation-order runs, the LLM-based method rankings remained highly concordant, with median pairwise Spearman correlations ranging from 0.876 to 0.984 and tie-corrected Kendall’s coefficients of concordance ranging from 0.866 to 0.984 across the six datasets (Supplementary Figure S10). The mean dataset-level difference in ARI between the randomized-order runs and the default ordering was −0.0481, with a 95% confidence interval of −0.1201 to 0.0240 and a *p*-value of 0.1469, indicating that no significant average shift in downstream performance was detected. Thus, although the default pipeline uses a fixed presentation orientation rather than reciprocal comparisons, the aggregate method rankings and downstream performance were relatively robust to presentation-order variation under the evaluated settings.

Evidence-accumulation clustering weights all selected partitions equally, so correlated methods may receive greater collective influence and reinforce shared errors. We therefore quantified redundancy using the Jaccard similarity between the pairwise co-assignment sets of selected methods. Across all method pairs selected by L-STAR, the mean co-assignment similarity was 0.355, compared with 0.400 for the fixed-consensus approach, indicating that adaptive selection did not increase overall redundancy relative to a fixed ensemble (Supplementary Figure S11). Nevertheless, 53 of the 60 selected method pairs involved two GNN-based methods, confirming that the selected partitions are not independent and may share common modeling biases. Similarity among these GNN-based pairs varied substantially, with a mean of 0.365 and a range from 0.133 to 0.659, indicating that methods within the same broad family can still generate substantially different partitions. Despite this redundancy and the absence of an explicit diversity constraint, L-STAR achieved strong and robust empirical performance across the evaluated datasets, outperforming the fixed-consensus strategy and individual methods overall. This suggests that the current adaptive selection and consensus procedure can effectively integrate partially correlated partitions in the evaluated settings. However, shared biases among correlated methods may still be reinforced through evidence accumulation, and incorporating explicit diversity constraints or redundancy-aware weighting represents an important direction for future development.

Although the benchmark study primarily used in this work was published after GPT-5’s knowledge cutoff, the original spatial transcriptomics datasets included in the benchmark were published before the cutoff. The visualizations evaluated in this study were either obtained from the post-cutoff benchmark study or generated using our own pipeline. Therefore, the exact visualizations evaluated are unlikely to have appeared in the pre-cutoff literature. Nevertheless, GPT-5 may retain prior knowledge of the corresponding anatomical regions, which could contribute to its assessment of biologically plausible spatial structures. Therefore, the current evaluation cannot completely disentangle de novo visual reasoning from previously acquired anatomical knowledge, and this possibility should be considered when interpreting the results.

## Methods

### L-STAR

#### Input

L-STAR can be applied to individual ST samples generated by both spot-based and image-based platforms. Spot-based methods, such as 10x Visium, generate gene expression profiles for spatial spots, where each spot may consist of multiple cells. Image-based methods, such as 10x Xenium, generate gene expression profiles for individual cells. Before running L-STAR, multiple single spatial domain detection methods, such as GraphST and SpaGCN, need to be applied to the ST data. By default, L-STAR requires as input the spatial locations of cells or spots, as well as the spatial domain assignment information from each single spatial domain detection method. L-STAR then internally generates images visualizing the spatial domain assignments for each method, using Palo^15^ to optimize the color palette. Alternatively, users can generate the spatial domain visualization images themselves and provide both the images and the corresponding domain assignment information as input to L-STAR. In both cases, an optional H&E image can be provided.

#### Step 1: Pairwise comparisons of single methods

L-STAR begins by conducting pairwise comparisons among single spatial domain detection methods using images that visualize their spatial domain assignments. Given results from *m* single methods and *R* repetitions (*R* = 5 by default), L-STAR evaluates each method pair sequentially, resulting in 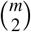 comparisons per repetition and 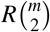 LLM calls in total. By default, L-STAR uses GPT-5 (version `gpt-5-2025-08-07`) with `temperature` = 1.0 and medium `reasoning effort`. Alternative LLMs are also supported, including Claude Sonnet 4 (version `claude-sonnet-4-20250514`) and Gemini 3 Pro (version `gemini-3-pro-preview`). Note that all LLMs evaluated in this study have knowledge cutoffs no later than January 2025.

In the default L-STAR pipeline, method names are sorted alphabetically before all unordered method pairs are enumerated, and within each pair the image of the alphabetically earlier method is always presented first. The method names themselves are not provided to the LLM, which receives the two candidate visualizations only as the first and second models. Each pair is therefore evaluated in a single fixed orientation per repetition, and reciprocal comparisons are not performed by default. To evaluate sensitivity to presentation order, we additionally performed 30 independent random-order runs for each of the six datasets. For every unordered method pair and each repeated comparison, assignment of the two method images to the first and second positions was randomized with equal probability. Across the 30 rounds, both presentation orientations occurred for every candidate pair. This analysis was performed as a sensitivity analysis and did not alter the default L-STAR pipeline.

For each pairwise comparison, the LLM is provided with two images visualizing the spatial domains identified by the two single methods under comparison, along with an optional H&E image when available. In addition, the LLM receives the following system-level and user-level prompts:

### System-level prompt

~~~
You are an expert model evaluator for spatial transcriptomics layer identification. Always start with EXACTLY one word: ‘first’ or ‘second’, then provide two short paragraphs in the form: First Model: reasoning; Second Model: reasoning
~~~

### User-level prompt

~~~
The slices belong to <DATASET_NAME>. Based on the information, please compare the model performance of
 identifying the layers of the slice provided in the next few messages.
[Attachment: H&E Image (optional)]
These two pictures are two identification results from two different models on this slice. Please compare which model performed better. Start your answer with EXACTLY ONE WORD: either ‘first’ or ‘second’; then give a brief but structured justification in one paragraph for each model in the format of ‘First Model: [reasoning] Second Model: [reasoning]’, highlighting the reasons for your choice.
[Attachment: Image 1 for the First Model]
[Attachment: Image 2 for the Second Model]
~~~

#### Step 2: Selection of top-performing methods

After completing all pairwise comparisons, L-STAR computes a winning rate for each method. For a given spatial domain detection method, the winning rate is defined as the proportion of pairwise comparisons in which the method is preferred by the LLM over its competitor, considering only comparisons involving that method. The top *k* methods with the highest winning rates are then selected, with *k* = 5 by default.

#### Step 3: Consensus clustering

Finally, L-STAR applies evidence-accumulation clustering (EAC)^18^ to integrate the spatial domain assignments from the *k* selected methods into a single consensus output. By accumulating co-assignment evidence across methods, EAC leverages complementary information from multiple high-performing candidates and produces a robust consensus set of spatial domain assignments.

Let *n* denote the number of spots or cells in the ST sample, and let *ℳ*_*k*_ denote the set of top-*k* methods selected in Step 2.

For the *ℓ*th method, let 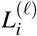 denote the spatial domain label assigned to the *i*th spot or cell.

L-STAR constructs an EAC matrix **C** *∈* [0, 1] defined as

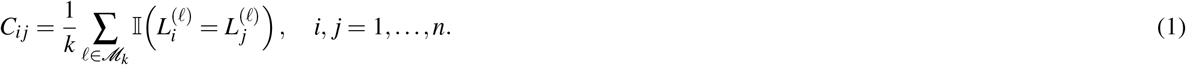

Here, *C*_*i j*_ quantifies the proportion of selected methods that co-assign spots *i* and *j* to the same spatial domain, with larger values indicating stronger agreement across methods.

L-STAR applies hierarchical clustering with average linkage to the distance matrix **D** = **1** − **C**, where **1** denotes the *n × n* matrix of ones. By default, the number of clusters is set to the median of the numbers of clusters produced by the selected top-*k* methods. Users may optionally specify an arbitrary number of clusters.

#### Optional cost-effective mode

L-STAR additionally provides an optional cost-effective mode that replaces the exhaustive pairwise comparisons in Step 1 with a single all-wise comparison. In this mode, visualizations from all *m* candidate methods are presented to the LLM simultaneously, each with a unique label, together with an optional H&E image when available. The LLM is then asked to return a strict ranking of all methods from best to worst. This reduces the number of LLM calls from 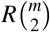 to *R*, where *R* denotes the number of repetitions. The system-level and user-level prompts retain the same overall structure as those used for pairwise comparisons, but instruct the LLM to return a JSON object containing a strict ranking in which every label appears exactly once and ties are not permitted. The complete prompts are provided in Supplementary Note 1. The same GPT-5 version and temperature are used as in the default mode, with high `reasoning effort`, because simultaneously ranking all *m* candidate methods requires more complex reasoning than selecting between two methods.

Accordingly, the method-ranking procedure in Step 2 is replaced by a rank-based scoring scheme. In each repetition, a method ranked *q*th receives a score of *m − q*, such that the highest-ranked method receives a score of *m −* 1, while the lowest-ranked method receives a score of 0. These scores are averaged across repetitions, and the top *k* methods with the highest average scores are selected for the consensus clustering procedure in Step 3, which remains unchanged.

#### Post-hoc spatial domain annotation

L-STAR includes an optional post-hoc module that assigns a biological name to each consensus domain without modifying the underlying domain assignments. The module takes as input the L-STAR spatial-domain assignments, gene expression data, and a user-declared sampling level of either spot or cell, which determines the granularity at which domains are annotated. Users may also provide optional dataset context, such as species, anatomical region, or tissue type, together with the expected scope of the annotation. Preprocessing, quality control, and marker identification are performed using `scanpy`^19^ with default hyperparameters. Raw-count inputs are normalized using `scanpy.pp.normalize_total` followed by `scanpy.pp.log1p`, whereas inputs that are already log-transformed undergo no additional transformation. Genes detected in fewer than three cells or spots are removed before marker computation. Marker genes for each domain are identified using `scanpy.tl.rank_genes_groups` and filtered using the default prevalence criteria implemented in `scanpy.tl.filter_rank_genes_groups`. The top 25 marker genes for each domain are retained for downstream annotation.

Annotation is performed once per dataset using GPT-5 (version `gpt-5-2025-08-07`) with high `reasoning effort`. GPT-5 receives information of all domains with their ranked marker genes, dataset context, sampling level, the L-STAR spatial-domain visualization, and an H&E image when available, and returns one biological name per domain. The prompt instructs the model to use standard anatomical or ontology nomenclature at the granularity supported by the markers, treating mitochondrial, ribosomal, and ubiquitously expressed genes as indicators of cell state rather than regional identity. Spot-level domains are named as regions, layers, or niches, whereas cell-level domains are named as cell types or states. Domains lacking sufficient marker evidence are assigned “Unknown”. The complete prompt is provided in Supplementary Note 2, and the output retains the observation identifier, L-STAR domain, assigned name, marker statistics, and structured model response for auditing.

### Performance evaluation

The clustering performance is evaluated using the adjusted Rand index (ARI), normalized mutual information (NMI), and adjusted mutual information (AMI). ARI values are computed using the `adjustedRandIndex` function from the `mclust` package in R. NMI and AMI are computed using the `NMI` and `AMI` functions from the `aricode` package in R. To ensure comparability across datasets, we apply a within-dataset linear normalization to the raw ARI, NMI, or AMI scores, mapping the worst-performing method to a normalized value of 0 and the best-performing method to a normalized value of 1.

### Color palette perturbation

Sensitivity to the color palette was evaluated by regenerating the visualizations of all candidate methods in each dataset under six transformations, comprising five random hue shifts and one grayscale conversion. Each hue shift applied a single rotation angle, drawn uniformly from 0 to 360 degrees, to the visualizations of all candidate methods within a dataset, preserving the spatial-domain assignments and spatial coordinates while changing only their chromatic representation. The grayscale images were obtained by replacing each pixel with its luminance, using weights of 0.299, 0.587, and 0.114 for the red, green, and blue channels, respectively. H&E images, when available, were left unmodified. The complete L-STAR pipeline was rerun under each condition, yielding 36 dataset-by-palette comparisons against the default palette. Agreement with the default condition was quantified by Kendall’s *τ*_*b*_ between the complete method rankings derived from the winning rates, by the Jaccard index between the selected top-*k* method sets with ties retained at the cutoff, and by Spearman’s *ρ* between the winning rates and the raw ARI values of the individual methods. Downstream spatial domain detection performance was compared using within-dataset normalization bounds defined by the original comparison set, which comprised the individual methods, L-STAR with the default palette, and the fixed consensus. Because L-STAR results obtained with transformed palettes were not used to redefine these bounds, differences relative to the default palette reflected changes in the LLM-based assessment independently of the normalization scale.

### Domain annotation evaluation

Manual annotations were first curated from the original dataset publications^20–24^. The mouse hippocampus dataset was excluded because the original publication provided only numeric cluster identifiers rather than biologically meaningful domain annotations. The curated manual annotations and corresponding L-STAR annotations are provided in Supplementary Table 1. For each spatial domain, both annotations were converted to vector embeddings using `BioLORD-2023`^25^, and their agreement was quantified by cosine similarity. To establish a null distribution, the manual annotations were permuted among domains within each dataset and the cosine similarities were recomputed. This permutation procedure was repeated 10,000 times for each dataset.

### API cost and runtime evaluation

Computational cost was evaluated by performing one representative pairwise comparison and one representative all-wise comparison for each of the six datasets. For each API call, wall-clock latency and token usage were recorded. API costs were calculated based on the OpenAI Platform pricing for `gpt-5-2025-08-07`.

### Evaluation datasets

Methods were evaluated on six ST datasets spanning diverse technologies and tissue types, including a human dorsolateral prefrontal cortex (DLPFC) section, generated using the 10x Genomics Visium platform; an E8.5 mouse embryo section profiled by Slide-seqV2; the SpatialMouseAtlas2020 mouse embryo dataset (embryo 1) generated using SeqFISH; a mouse hippocampus section from the Spatial Transcriptomics (ST) platform; a mouse visual cortex (MVC) section profiled by STARmap; and an axolotl brain (AB) section generated using Stereo-seq.

The spatial coordinates of spots or cells and the corresponding spatial domain annotations produced by single spatial domain detection methods were obtained from the benchmarking study^9^. For the DLPFC dataset, the Slide-seqV2 mouse embryo section, and the SeqFISH SpatialMouseAtlas2020 dataset, spatial domain visualizations were cropped directly from figures in the benchmarking study^9^ and used as inputs to L-STAR. For the remaining three datasets, the default L-STAR input pipeline was applied.

The spatial-domain assignments produced by the individual methods were taken directly from the outputs released by the benchmarking study^9^ and were not independently regenerated, reclustered, or retuned. Accordingly, the same clustering algorithm was not imposed across methods. Each method retained the clustering procedure, preprocessing strategy, and parameter settings used in the original benchmark. The benchmark study reported that default parameter settings and preprocessing procedures were used unless otherwise specified. For methods requiring a predefined number of clusters, the number was set according to the annotated categories, whereas for methods parameterized by clustering resolution, the resolution was selected to produce a number of spatial domains as close as possible to the annotated number. The annotated numbers of clusters for the individual datasets are reported in Supplementary Table S1 of the benchmark study^9^, and additional details on clustering algorithms, preprocessing procedures, and default or customized parameter settings, including those for the GNN-based methods, are provided in its Methods and Supplementary Table S4^9^.

### Competing methods

The spatial domain assignments of 12 single spatial domain detection methods were directly obtained from the benchmarking study^9^. These methods include BayesSpace^5^, CCST^26^, conST^27^, GraphST^6^, SCGDL^28^, SEDR^29^, Seurat^30^, SpaGCN^7^, SpaceFlow^31^, Spatial_MGCN^32^, STAGATE^33^, and STMGCN^34^.

The highest-ranked method selected by GPT-5 was defined as the method with the highest winning rate, following the same procedure as L-STAR but without performing consensus clustering.

In addition, the same EAC procedure used by L-STAR was applied to a fixed set of representative methods recommended by the benchmarking study^9^, namely GraphST, BayesSpace, SpaGCN, and STAGATE.

### Keyword extraction from LLM responses

For each dataset, the GPT-5 outputs of the pairwise comparison rationales were pooled across all pairwise comparisons and repeated runs into a dataset-specific corpus. The rationale text was extracted from the JSONL files automatically generated by L-STAR during the pairwise comparison step and converted into terms using a `spaCy` pipeline with tokenization and lemmatization, where a small set of token-normalization rules was manually specified to maintain biological interpretability (e.g., normalizing “wm” as “white_matter”, and capturing “white matter” as the phrase “white_matter”). Each alphabetic token was assigned a part-of-speech tag, and terms were defined based on content-bearing tokens, including noun or proper-noun unigrams and adjective-noun phrases. Uninformative terms with high occurrence proportion were removed automatically by combining standard English stopwords from `sklearn` and `WordCloud` libraries, together with dataset-specific stopword lists curated from a prior TF-IDF analysis and implemented as fixed global stopword lists, including articles (e.g., “a”, “an”, “the”), decision-template tokens (e.g., “first”, “second”, “both”), generic pipeline and comparison nouns (e.g., “model”, “prediction”, “labels”), ubiquitous layout or spatial narration terms (e.g., “left”, “layer”, “region”), discourse scaffolding verbs (e.g., “indicate”, “suggest”, “show”), subjective stance/evaluation words (e.g., “overall”, “better”, “poor”), and common artifact descriptors (e.g., “noise”). In addition to the automatically generated stopword lists, dataset-specific whitelists were manually specified by examining the dataset-specific TF-IDF results and selecting terms that are biologically informative. The occurrence proportion of a term was defined within each dataset as its pooled count divided by the total number of all terms after filtering in all pairwise comparisons and repeated runs.

## Acknowledgments

The study was supported by the National Institutes of Health under Award Number R35GM154865, U54AG075936, and U54AI191253.

## Author contributions

Z.J. conceived the study. C.Z. conducted the analyses and developed the L-STAR method. C.Z. and Z.J. wrote the manuscript.

## Competing interests

All authors declare no competing interests.

## Data availability

The spatial-domain assignments produced by the individual methods and analysed in this study were provided directly by the authors of the benchmarking study by Kang et al.^9^. The corresponding preprocessed spatial transcriptomics datasets are publicly available from the Benchmark ST analysis collection on Figshare: human dorsolateral prefrontal cortex section 151673 profiled by 10x Genomics Visium (https://doi.org/10.6084/m9.figshare.28200299.v1); an E8.5 mouse embryo profiled by Slide-seqV2 (https://doi.org/10.6084/m9.figshare.28200323.v1); SpatialMouseAtlas2020 embryo 1 profiled by seqFISH (https://doi.org/10.6084/m9.figshare.28195694.v1); mouse hippocampus section 2 profiled using the Spatial Transcriptomics platform (https://doi.org/10.6084/m9.figshare.28195169.v1); mouse visual cortex profiled by STARmap (https://doi.org/10.6084/m9.figshare.28195031.v1); and axolotl brain profiled by Stereo-seq (https://doi.org/10.6084/m9.figshare.28200305.v1).

## Code availability

The L-STAR software package is freely available at https://github.com/Williamzcy0929/L-STAR.

## Notes

### Competing Interest Statement

The authors have declared no competing interest.

