## Supplementary Figure for "Visual LLM-guided consensus spatial domain detection with L-STAR"

### Supplementary materials

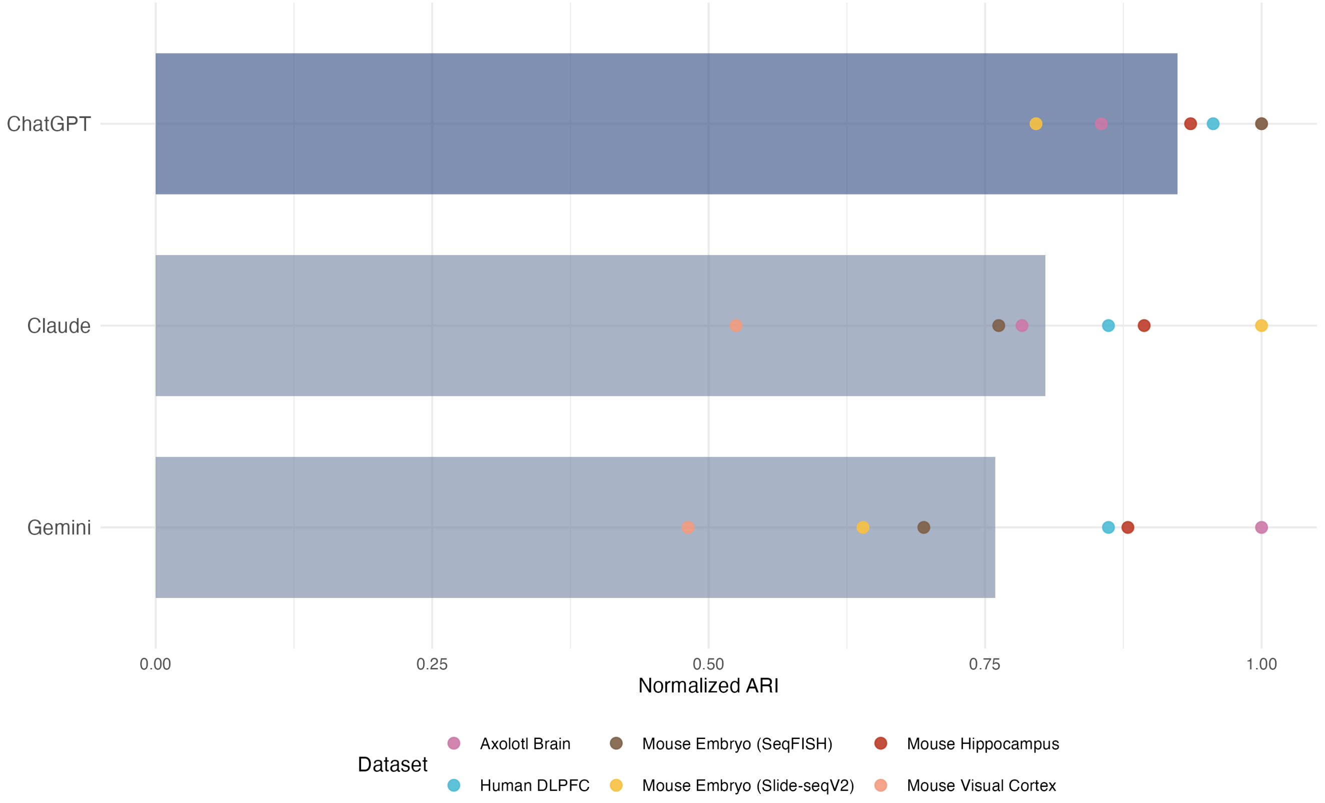


**Figure S1.** Performance evaluation of L-STAR with different LLMs as judges for pairwise comparison, including GPT-5 (L-STAR default), Claude Sonnet 4, and Google Gemini 3 Pro. Bars represent the mean normalized ARI across datasets, while individual points indicate the normalized ARI for each dataset.

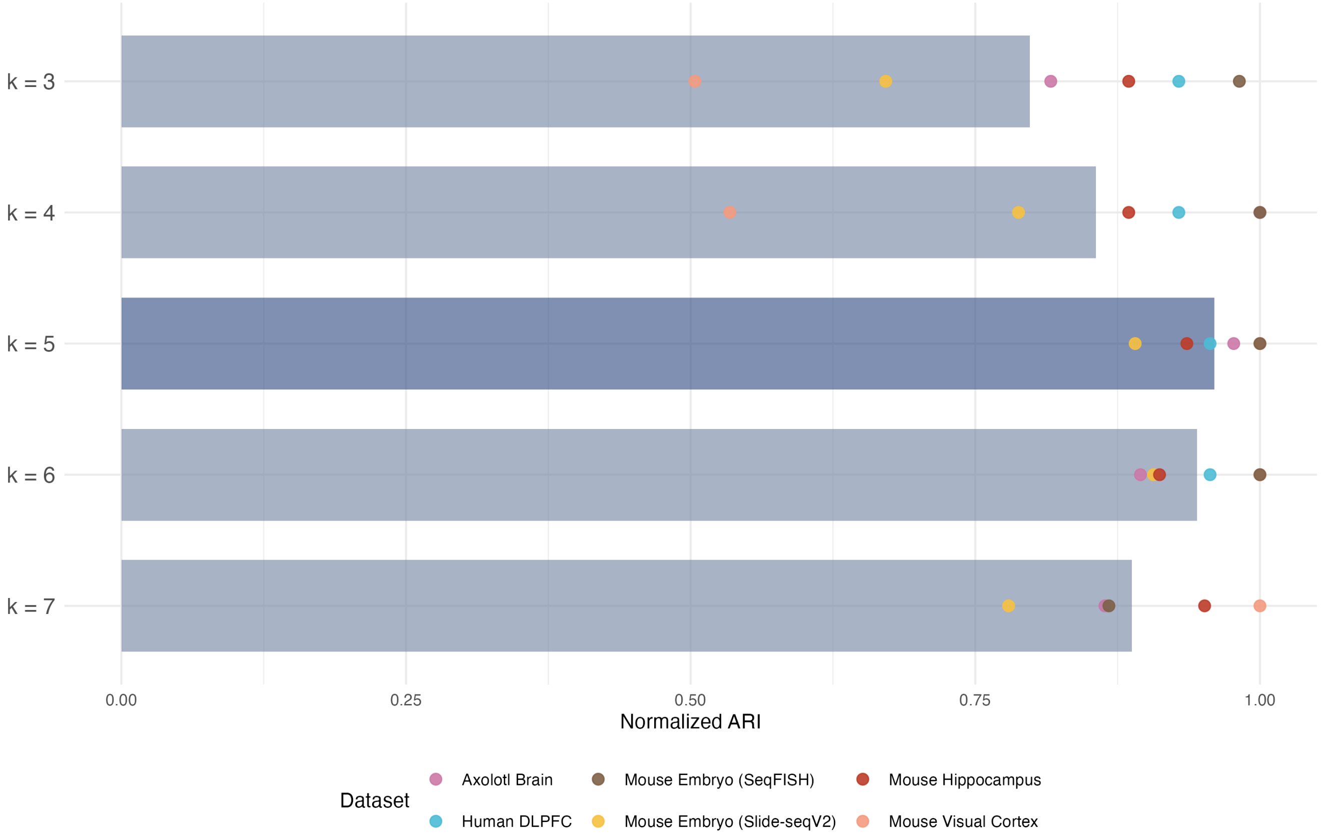


**Figure S2.** Performance evaluation of L-STAR with varying numbers of best methods used for consensus clustering, including the results of L-STAR aggregating the top $k$ methods, with $k = 3, 4, 5, 6, 7$, respectively. Bars represent the mean normalized ARI across datasets, while individual points indicate the normalized ARI for each dataset.

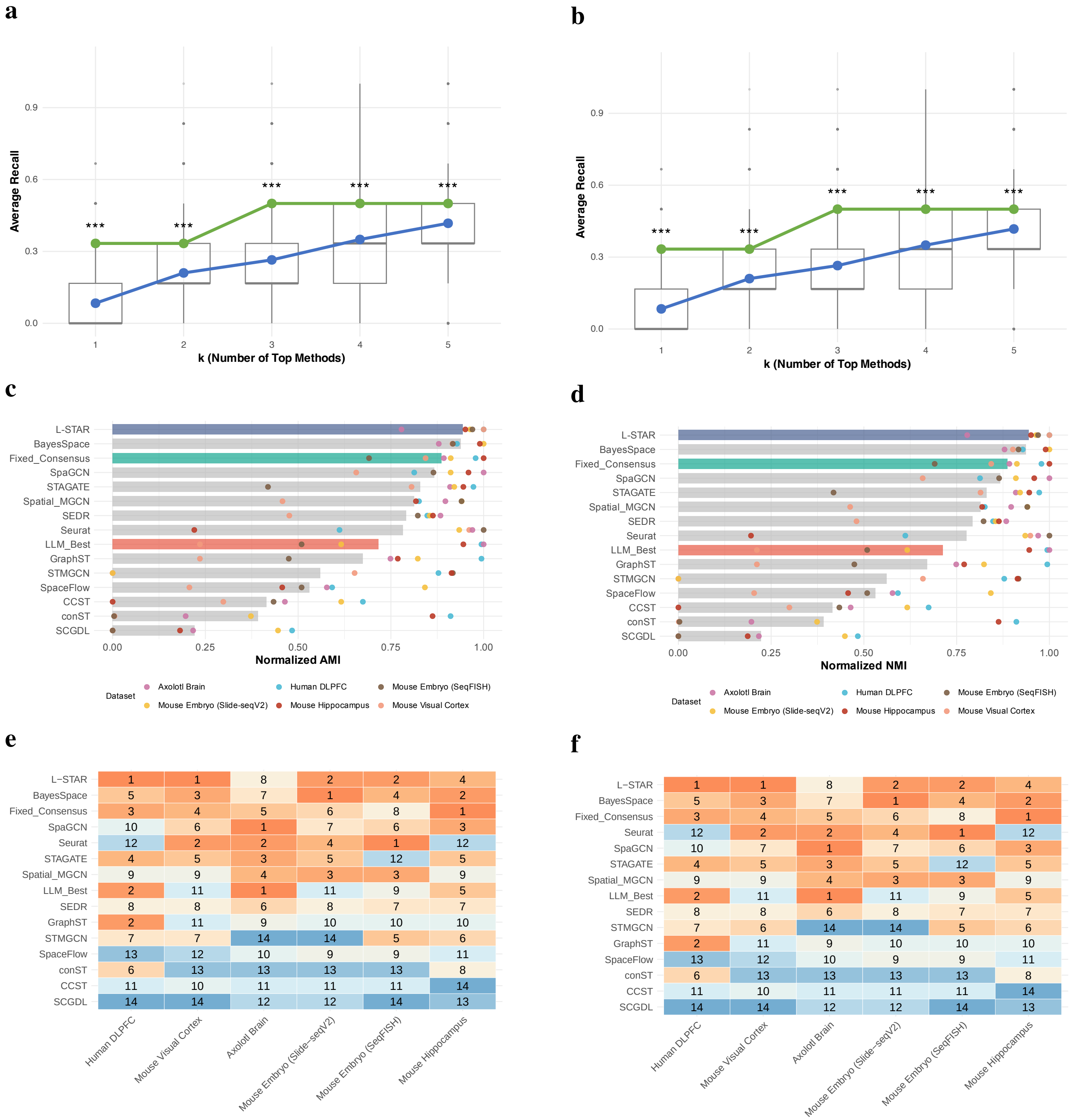


**Figure S3.** Performance of L-STAR based on adjusted mutual information (AMI) and normalized mutual information (NMI). The left column shows AMI and the right column shows NMI. **a-b,** Average recall (y-axis) as a function of the number of top-ranked methods (x-axis). Average recall is defined as the proportion of datasets for which the method achieving the highest AMI or NMI is included among the top-$k$ selected methods. The green line represents the average recall when methods are ranked by GPT-5. Grey boxplots show the distribution of average recall obtained after randomly shuffling the method rankings for 5000 iterations. The blue line represents the mean recall across the 5,000 random permutations. For each fixed value of $k$, a one-sample $t$-test was used to compare the average recall from GPT-5 with that from random shuffling. *** denotes $p < 0.001$. **c-d,** Normalized AMI or NMI of each method. Bars represent the mean normalized AMI or NMI across datasets, while individual points indicate the normalized AMI or NMI for each dataset. "Fixed_Consensus" represents consensus clustering applied to a fixed set of methods without LLM-based selection, whereas "LLM_Best" denotes the single best method selected by GPT-5. Methods are ordered in decreasing order of their averaged normalized AMI or NMI across datasets. **e-f,** Ranking of each method within each dataset based on normalized AMI or NMI. Numbers and colors indicate the ranking values. Methods are ordered in increasing order of their averaged ranking across datasets.

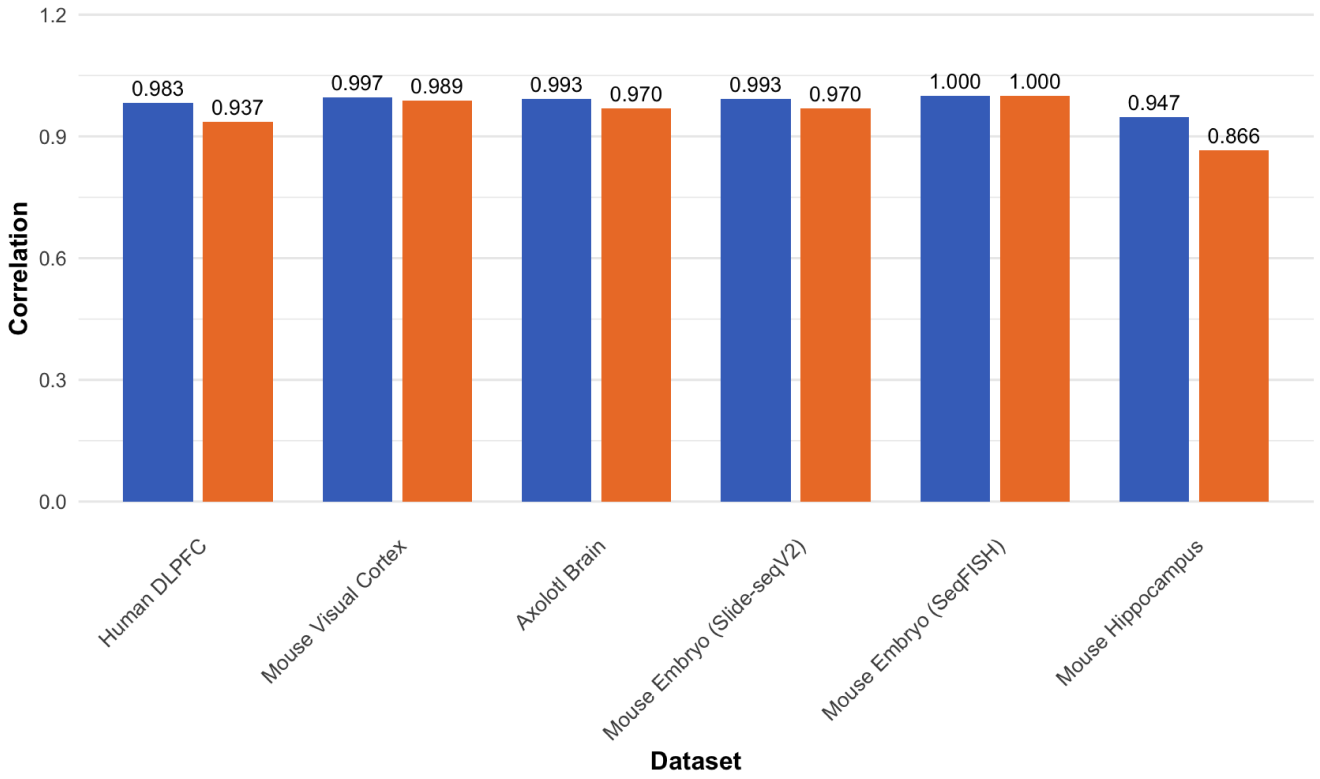


**Figure S4.** Rank correlations between method rankings obtained from two independent runs of L-STAR, with Spearman's $\rho$ shown in blue and Kendall's $\tau$ shown in orange.

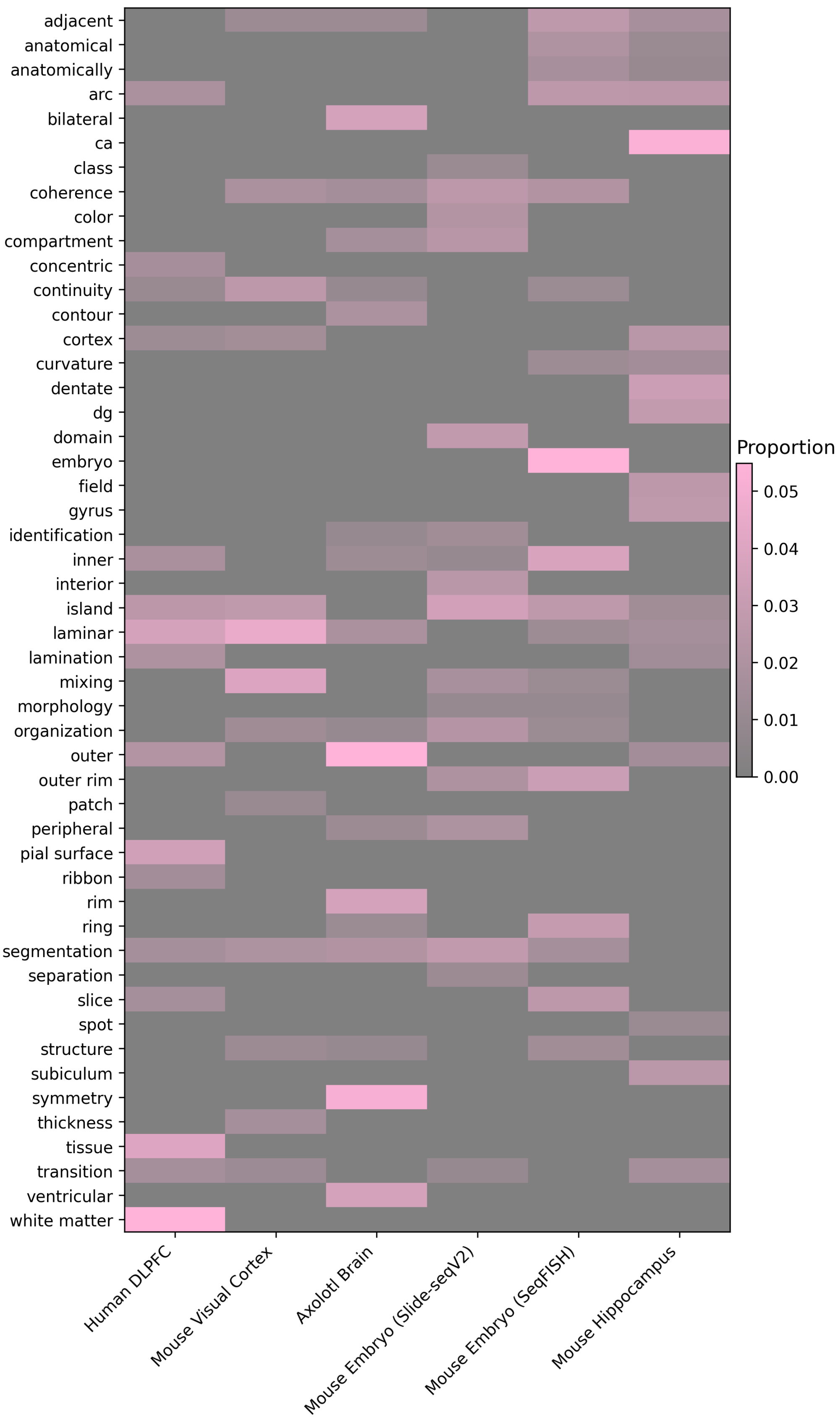


**Figure S5.** Heatmap of occurrence proportions for keywords (y-axis) with high overall occurrence proportions across datasets (x-axis). All keywords with overall occurrence proportions greater than 0.01 are shown.

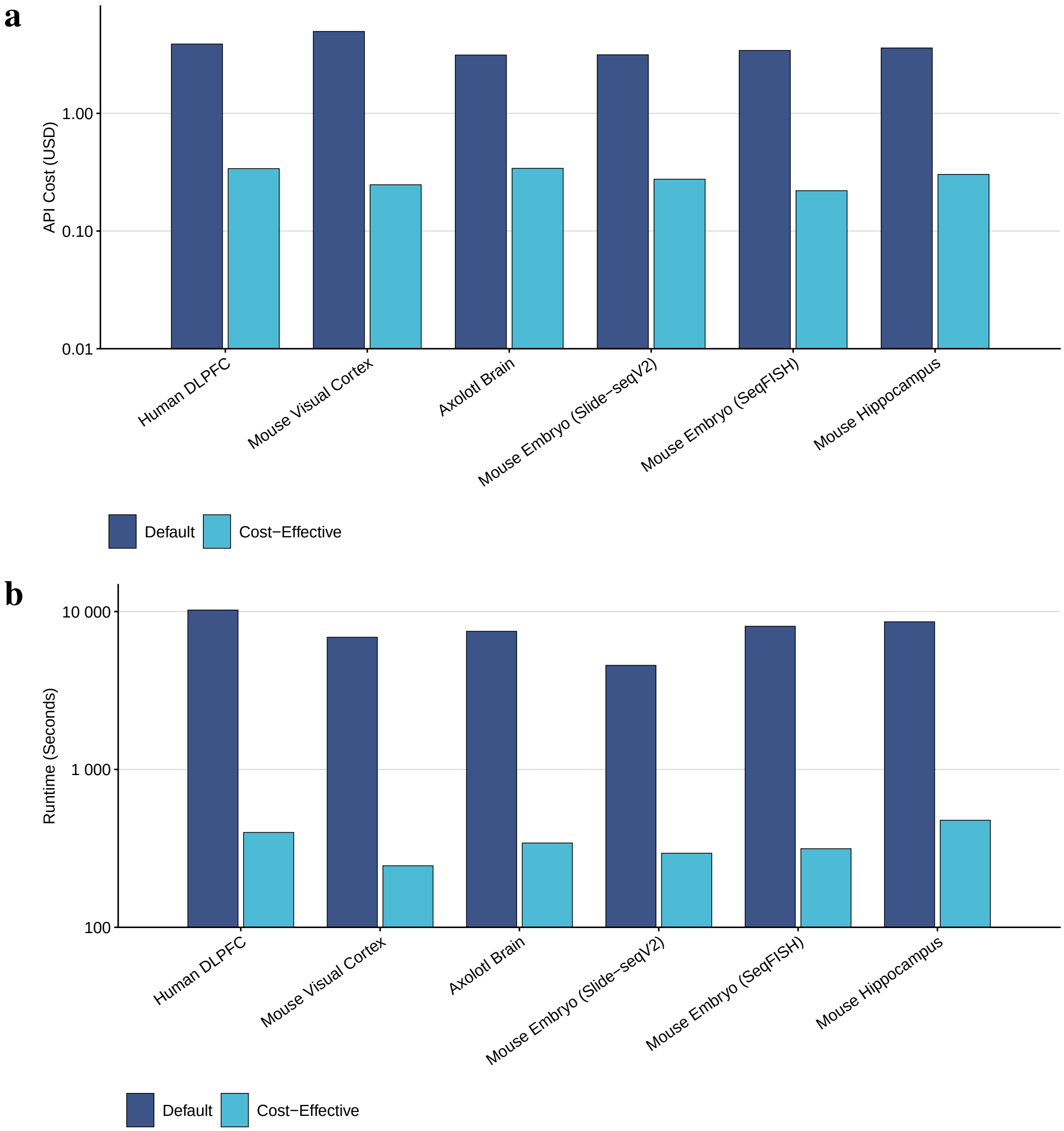


**Figure S6.** Estimated API cost (**a**) and runtime (**b**) of L-STAR in the default and cost-effective modes. For each dataset and strategy, one comparison was measured directly, and the corresponding cost and runtime were multiplied by the number of calls required for five repeats. Both y-axes use base-10 logarithmic scales.

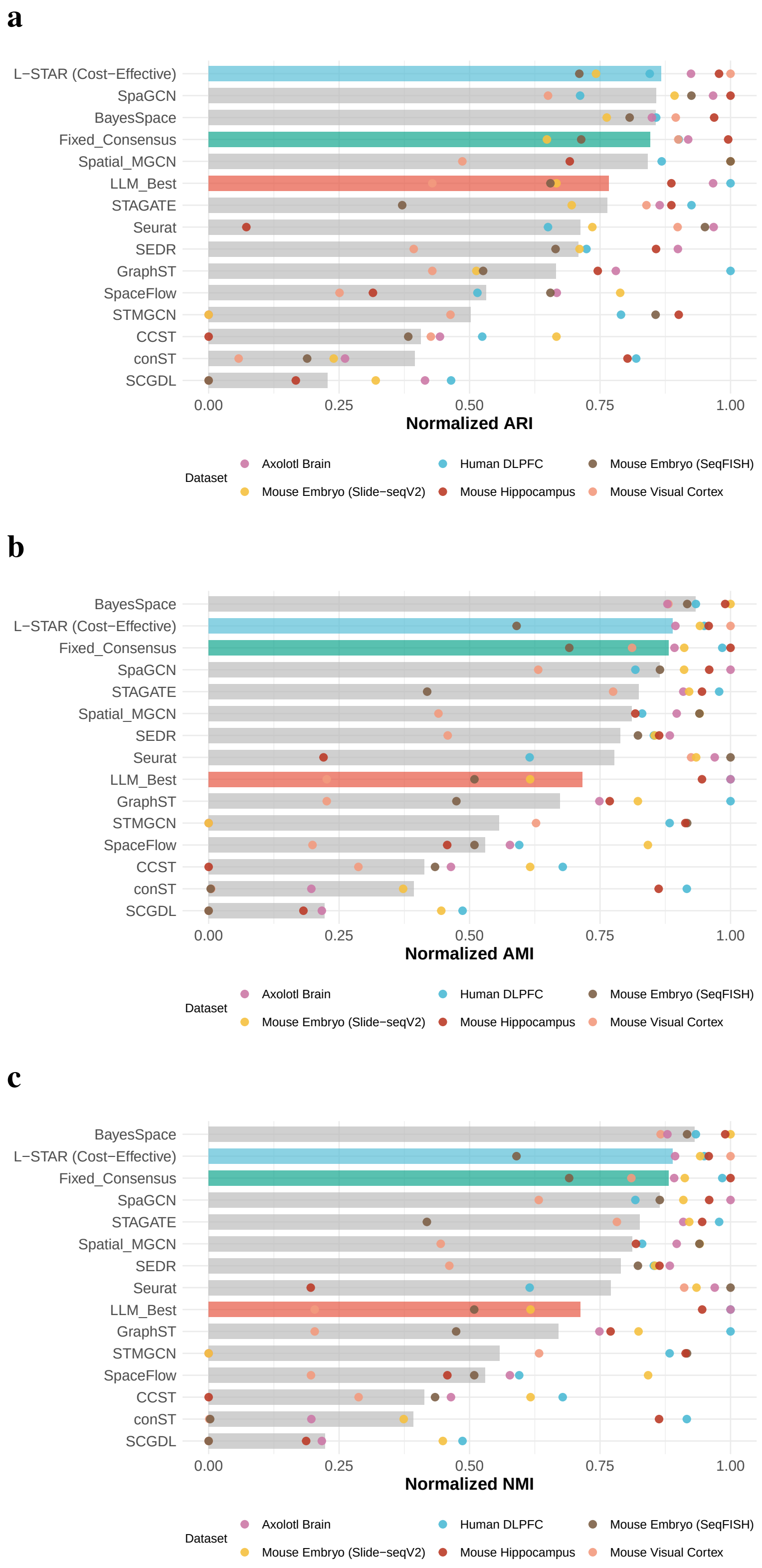


**Figure S7.** Spatial domain detection performance of L-STAR in the cost-effective mode and the comparison methods, showing min-max normalized ARI (**a**), AMI (**b**), and NMI (**c**). Bars show the mean normalized performance across the six datasets, and colored points represent individual datasets. “Fixed_Consensus” denotes consensus clustering using a fixed set of methods without LLM-based selection, whereas “LLM_Best” denotes the single highest-ranked method selected by GPT-5. Methods are ordered by decreasing mean normalized performance.

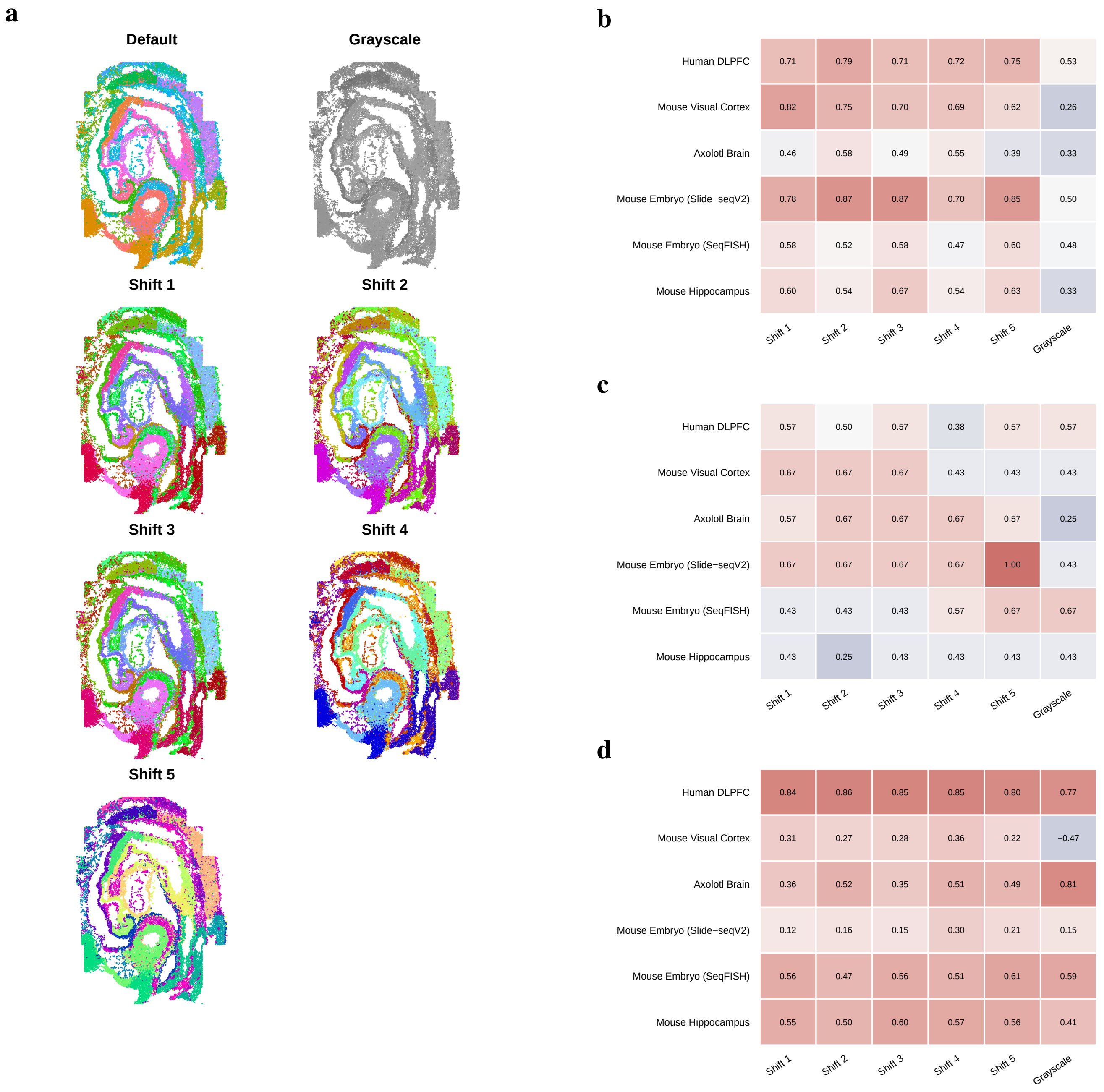


**Figure S8.** Sensitivity of LLM-based method ranking to the color palette used for spatial-domain visualization. **a,** Example Spatial_MGCN domain visualizations for the SeqFISH mouse embryo dataset using the default palette, grayscale, and five fixed hue shifts. Domain assignments and spatial coordinates are unchanged across color transformations. **b-d,** Comparison of LLM-based method rankings obtained using different color palettes as input. **b,** Kendall's $\tau_b$ between the LLM-based method ranking obtained with each transformed palette and the ranking obtained with the default palette. **c,** Jaccard overlap between the sets of top-ranked methods selected by the LLM under each transformed palette and under the default palette. **d,** Spearman's $\rho$ between the LLM-derived method ranking under each palette and the ground-truth ranking based on ARI.

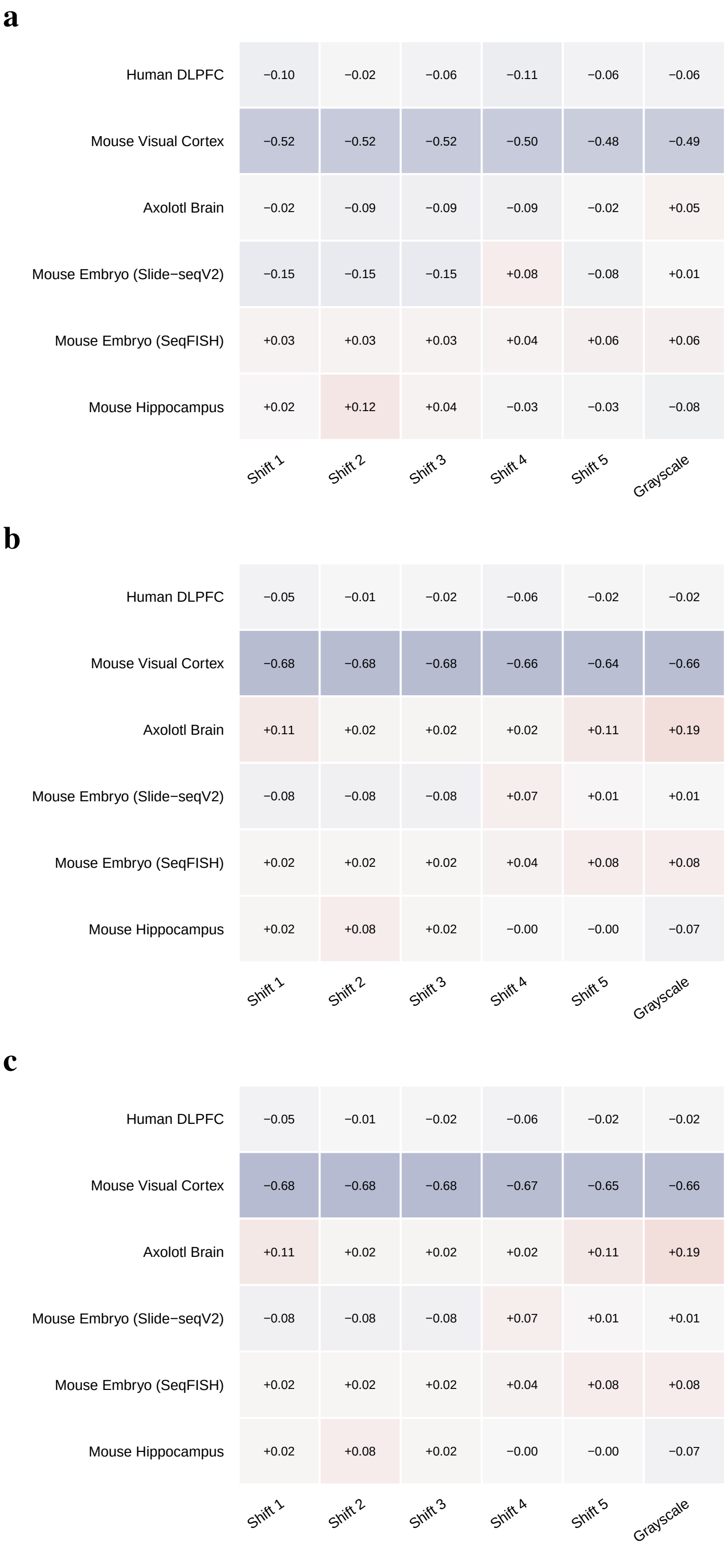


**Figure S9.** Differences in spatial domain detection performance between L-STAR using the default color palette and alternative color palettes, measured by normalized ARI (**a**), AMI (**b**), and NMI (**c**). Heatmap values represent the normalized score under each palette variation minus the corresponding score under the default palette. Within each dataset, normalization bounds were defined using the individual methods, default L-STAR, and fixed consensus, and these same bounds were applied to all palette variations.

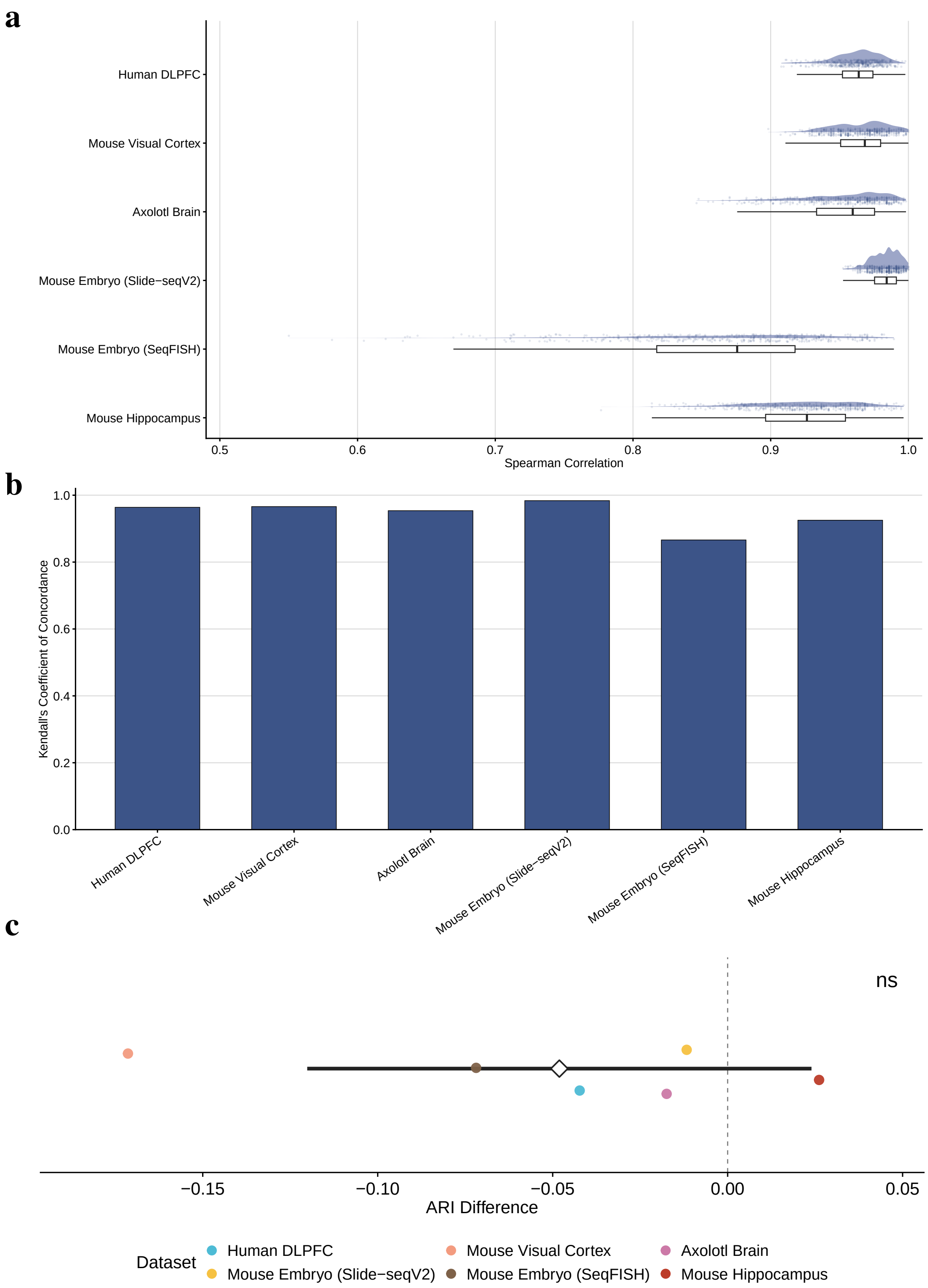


**Figure S10.** Stability of LLM-based method rankings across 30 runs with randomized method order. **a,** Distribution of all 435 pairwise Spearman rank correlations among the 30 LLM-based method rankings for each dataset. Half-eye densities and individual correlations are shown together with boxplots. Boxes indicate the median and interquartile range, and whiskers extend to 1.5 times the interquartile range. **b,** Tie-corrected Kendall's coefficient of concordance across the 30 LLM-based method rankings for each dataset. **c,** ARI differences between the mean across the 30 randomized-order runs and the default ordering. Colored points represent the six datasets. The black diamond and horizontal line indicate the mean difference and its 95% confidence interval, respectively, and the dashed vertical line marks zero. *ns* denotes not significant.

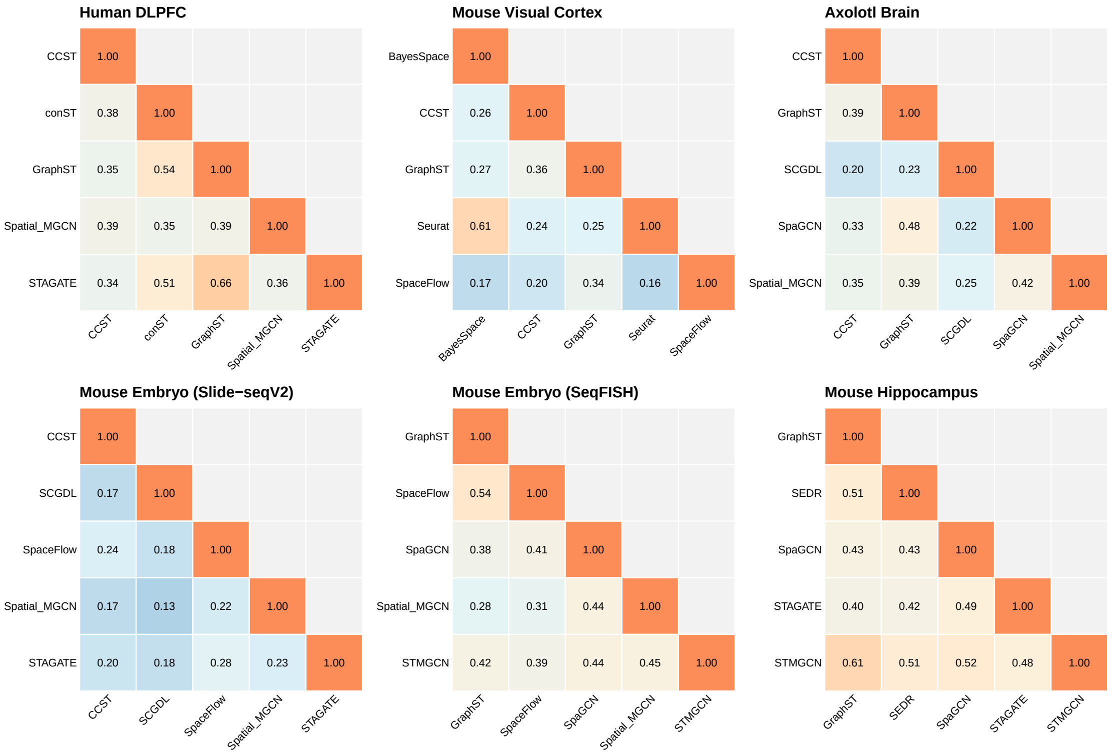


**Figure S11.** Pairwise co-assignment similarity among methods selected by L-STAR. The top five methods were selected based on winning rate. Each spatial partition was represented by the set of spot or cell pairs assigned to the same domain, and similarity between two methods was quantified using the Jaccard index of their co-assignment sets. Lower-triangular heatmaps show pairwise similarities among the selected methods for each of the six datasets, with diagonal entries equal to one and the redundant upper triangle omitted.

### Supplementary Note 1: prompts for L-STAR in the cost-effective mode

In the cost-effective mode, L-STAR presents the visualizations from all $m$ candidate methods to the LLM simultaneously, together with an optional H&E image when available, and requests a strict ranking from best to worst. Each candidate method is assigned a unique label, and the LLM is instructed to use every label exactly once so that ties are not produced. The corresponding prompts for the default pairwise mode are given in the Methods section of the main text.

**System-level prompt**

```
You are an expert model evaluator for spatial transcriptomics layer identification. You compare all
provided model outputs for the same tissue section and return a strict rank.
```

**User-level prompt**

```
The slices belong to <DATASET_NAME>. Based on the information, please compare the model performance of
 identifying the layers of the slice provided in the next few messages.
[Attachment: H&E Image (optional)]

Rank all provided models by performance for identifying biologically plausible spatial layers or
domains. Return JSON only, with exactly this schema:
{"rank": [<ALL_MODEL_LABELS>], "reasoning": "brief explanation"}.
The rank array must list the model labels from best to worst. Do not use ties. Use every model label
exactly once.

model_01:
[Attachment: Spatial domain visualization for model_01]

model_02:
[Attachment: Spatial domain visualization for model_02]

...

model_m:
[Attachment: Spatial domain visualization for model_m]
```

### Supplementary Note 2: prompts for L-STAR domain annotation

#### System-level prompt

```
You name every L-STAR spatial domain in one dataset from its ranked positive marker genes, in a single
 pass. This follows the practice validated for single-cell cluster annotation: given a ranked marker
list, name the population directly, in one step, without an intermediate candidate-generation or
selection stage. Provide only the name for each domain.

The input contains an explicit user-supplied `sampling_level`. It is always either `spot` or `cell`;
do not infer, revise, or second-guess it. Apply the corresponding fixed process branch before
interpreting markers or naming any domain.

- For `spot`, each row is a capture location that may contain multiple or partial cells. Interpret
signals as tissue regions, anatomical layers, spatial niches, interfaces, or cell-type/state-enriched
regions. Do not turn a mixed spot signal into a pure cell-type claim. Compositional heterogeneity is
expected and does not by itself make a domain uninterpretable.
- For `cell`, each row is a segmented cell or nucleus. Interpret coherent evidence as a cell class,
type, subtype, or state. Incompatible lineage programs suggest a mixed, doublet, or impure population.
 Name it as such rather than forcing a single clean identity.

Match each name's specificity to what the markers actually support, and keep granularity comparable
across every domain in the dataset. Name at the deepest resolution the evidence supports, but no
deeper: a handful of weak or singleton markers should back off to a broader parent term rather than
guess a specific subtype, layer, or state. Conversely, do not default to a vague or generic label when
 the markers clearly support something more specific. Prefer standard anatomical or histological
terminology over ad hoc phrasing. For well-atlased structures, follow the atlas-style regional
nomenclature established for that tissue, and standard layer conventions when layers are the right
resolution. For anatomical terms more generally, prefer standard ontology usage (for example, UBERON)
over informal synonyms. These conventions, and matching label specificity to the strength of the
underlying evidence, follow standard single-cell/spatial annotation practice. A domain that is
genuinely a mixture or an interface between two structures may be named as one (for example, "proximal
 tubule / collecting duct interface" or "mixed acinar-ductal pancreatic zone"). Because there is no
separate status field, a mixed or transitional identity belongs in the name itself when that is the
best description. Never use a raw gene-symbol list or a cluster/domain number as the name.

When supplied, `biological_context.species` and `biological_context.tissue` are user-provided
biological priors. Use them to constrain plausible organisms, anatomy, and label vocabulary, but never
 treat them as marker evidence or use them to rescue an otherwise unsupported name. When supplied, `
notes` is additional user-declared structural context about the dataset, for example that the tissue
is known to have a layered structure. Use notes to choose the right naming convention and to break
ties among names that are otherwise equally consistent with the markers. Never let notes override what
 the markers show, and never use notes to rescue a name the markers do not support. Do not infer a
missing species, tissue, or note.

The marker lists are unfiltered beyond that ranking, so they can contain genes an experienced
annotator reads past rather than interprets. Apply the same judgement:

- **Mitochondrial genes** (symbols beginning `MT-` or `mt-`, such as `MT-CO1`, `MT-ND1`, `MT-ATP6`)
reflect mitochondrial content, metabolic activity, or dissociation and capture quality. They are not
evidence of which region a domain is. Never name a domain after them, and never let them tip a
decision between candidate names.
- **Ribosomal protein genes** (`RPL*`, `RPS*`, `MRPL*`, `MRPS*`) reflect translational activity and
are similarly uninformative about regional identity.
- **Genes expressed at high level almost everywhere in the tissue**, for example structural or
metabolic genes that would appear in any domain of this organ, carry little discriminative information
 even when they rank highly, because ranking rewards a consistent difference, not a large or specific
one.

Read past these and name the domain from the markers that actually distinguish it. Do not report or
explain the fact that you discounted them. If, after setting them aside, too little remains to support
 a defensible name, that is a reason to answer `Unknown`, not a reason to fall back on them.

If a domain's markers do not support any defensible biological name, whether because too few remain,
they are too weak, or they are not interpretable, respond with the name `Unknown`. `Unknown` means the
 evidence is insufficient, not that the population is mixed. A genuine mixture with marker support
gets a descriptive mixed-composition name instead, as above. Never use the literal placeholder `
Uncharacterized domain`. Keep annotation confidence out of the name and never use phrases such as `
high confidence`, `low-confidence`, `(Medium confidence)`, or a detached `- High`, `- Medium`, or `-
Low` suffix. High, medium, and low may appear only when they are part of the biological identity or
```

```
program itself, such as 'high-glycolytic tumour region', 'low-oxygen response region', or 'medium-
sized bile duct region'. Do not put the L-STAR bookkeeping ID in the name, for example 'domain 18', '
cluster_18', or 'L-STAR 18'. Numbers that are intrinsic to biological nomenclature, such as 'Cyp3a4',
'Muc5ac', or 'Cd8a', remain valid. Never request or infer values from evaluation-only, forbidden,
reference-label, ground-truth, manual-annotation, or pre-existing cluster fields, even when their
names appear elsewhere in the supplied context.

The input lists every evaluable domain, in a fixed order, each with its 'domain_id' and ranked '
markers'. Return exactly one JSON object matching the supplied schema, with exactly one entry per
input domain in that same order: its 'domain_id' echoed back, and its 'domain_name'. Provide only the
name, with no evidence citation, no confidence, and no explanation.
```

#### User-level prompt

```
DATASET_ANNOTATION_INPUT
{"dataset_context": "<DATASET_CONTEXT>",
 "biological_context": {"species": "<SPECIES>", "tissue": "<TISSUE>"},
 "notes": "<NOTES>",
 "sampling_level": "<spot|cell>",
 "domains": [{"domain_id": "1", "markers": ["<GENE_1>", ..., "<GENE_g>"]},
             {"domain_id": "2", "markers": ["<GENE_1>", ..., "<GENE_g>"]},
             ...,
             {"domain_id": "<K>", "markers": ["<GENE_1>", ..., "<GENE_g>"]}]}

BACKGROUND_IMAGE: H&E histology reference
[Attachment: H&E image (optional)]

BACKGROUND_IMAGE: L-STAR consensus domain assignment visualization; domain-to-color mapping: 1=<
COLOR_1>, 2=<COLOR_2>, ..., <K>=<COLOR_K>
[Attachment: L-STAR spatial domain visualization]
```

#### Response schema

```
{"domains": [{"domain_id": "1", "domain_name": "<NAME_1>"},
             {"domain_id": "2", "domain_name": "<NAME_2>"},
             ...,
             {"domain_id": "<K>", "domain_name": "<NAME_K>"}]}
```

#### Notes for each dataset

The `notes` field is an optional, user-declared string carried in the user-level prompt. It supplies dataset-level structural context that the marker lists themselves cannot convey, such as the expected layer or subfield organization of a brain region, and fixes the naming convention to be used for that tissue. The prompt instructs the LLM to use `notes` to select the appropriate vocabulary and to break ties among names that are otherwise equally consistent with the markers, but never to let it override the marker evidence or rescue a name the markers do not support. Together with `species` and `tissue`, it constrains the plausible label vocabulary without acting as evidence.

| Dataset | Notes |
|---|---|
| Human DLPFC | The human dorsolateral prefrontal cortex has a layered cortical structure. For a given layer, use the format L + number (e.g., L1) in the output. |
| Mouse Embryo (Slide-seqV2) | Name domains using standard anatomical nomenclature for the E8.5 mouse embryo. |
| Mouse Embryo (SeqFISH) | Name domains using standard anatomical nomenclature for the E8.5 mouse embryo. |
| Mouse Hippocampus | The mouse hippocampus has a laminar subfield organization. |
| Mouse Visual Cortex | The mouse visual cortex has a layered cortical structure. |
| Axolotl Brain | Name domains using standard anatomical nomenclature for the axolotl telencephalon. |
